# Mental imagery can generate and regulate acquired differential fear conditioned reactivity

**DOI:** 10.1101/2021.02.04.429795

**Authors:** Steven G. Greening, Tae-Ho Lee, Lauryn Burleigh, Laurent Grégoire, Tyler Robinson, Xinrui Jiang, Mara Mather, Jonas Kaplan

## Abstract

Mental imagery is an important tool in the cognitive control of emotion. The present study tests the prediction that visual imagery can generate and regulate differential fear conditioning via the activation and prioritization of stimulus representations in early visual cortices. We combined differential fear conditioning with manipulations of viewing and imagining basic visual stimuli in humans. We discovered that mental imagery of a fear-conditioned stimulus compared to imagery of a safe conditioned stimulus generated a significantly greater conditioned response as measured by self-reported fear, the skin conductance response, and right anterior insula activity (experiment 1). Moreover, mental imagery effectively down- and up-regulated the fear conditioned responses (experiment 2). Multivariate classification using the functional magnetic resonance imaging data from retinotopically defined early visual regions revealed significant decoding of the imagined stimuli in V2 and V3 (experiment 1) but significantly reduced decoding in these regions during imagery-based regulation (experiment 2). Together, the present findings indicate that mental imagery can generate and regulate a differential fear conditioned response via mechanisms of the depictive theory of imagery and the biased-competition theory of attention. These findings also highlight the potential importance of mental imagery in the manifestation and treatment of psychological illnesses.

## INTRODUCTION

From fears of monsters under our beds to remembering the ‘good times’ while dealing with the death of a loved one, mental imagery is central to the cognitive control of emotion[1–3]. Modern frameworks for conceptualizing the cognitive control of emotion – specifically emotion generation and emotion regulation – have similarly suggested the importance of mental imagery. Gross[4] suggests a process model in which internal situations (e.g., imagined objects or situations), in addition to external ones, can produce an emotional response. Likewise, mental imagery has been discussed as a potential factor in both distraction and reappraisal strategies for regulating emotions[3, 5].

Fear (or threat) conditioning is one of the most well-studied paradigms for investigating both the generation[6–8] and the regulation[9–14] of emotional reactivity. In differential fear conditioning a neutral conditioned stimulus (CS+) is paired with an aversive unconditioned stimulus (US), such as a mild shock, while a second conditioned stimulus (CS-) is never paired with the US. Following successful fear conditioning, presentation of the CS+ alone elicits a measurable conditioned response (CR, e.g., self-reported subjective fear; a skin conductance response, SCR; or differential neural activity) compared to the CS-.

While much research in human and non-human animals has emphasized the role of the amygdala in the acquisition of differential fear conditioning[15–22], a recent meta-analysis of functional magnetic resonance imaging (fMRI) studies found that a network that includes both cortical and subcortical regions but *not* the amygdala is reliably activated by differential fear conditioning[6]. This putative fear network includes bilateral aspects of the anterior insula (aIn), dorsal anterior cingulate cortex (dACC), and dorsomedial prefrontal cortex (dmPFC), ventral striatum, thalamus, and midbrain structures. Notably, the largest effect sizes of differential conditioning were observed in the left and right aIn, and the dmPFC, respectively[6], potentially reflecting the role of these regions in the expression of fear or the conscious anticipation of threat [6,23,24]. As the present study is primarily concerned with the expression and regulation of differential conditioning rather than the mechanisms of acquisition, we prioritized the aIn and dmPFC in the univariate brain analyses.

Mental imagery also appears to interact with differential fear conditioning [25, 26]. For example, Reddan et al.[27] found that mental imagery of the auditory CSs could be used to induce extinction learning following differential fear conditioning to auditory tones, as measured using SCR and whole-brain multivariate pattern analysis. In addition, mental imagery of CSs can contribute to extinction learning[28, 29] or facilitate the reconsolidation of differential fear conditioning[30]. In the current study, we evaluated whether visually acquired differential conditioning transfers to imagery of the CSs and whether the mechanisms for imagery generation are consistent with the depictive theory of mental imagery.

According to the depictive theory[31], mental imagery is the top-down production of neural representations from memory, which are similar to those produced by perception and is associated with a conscious experience of ‘seeing in the mind’s eye’[31, 32]. Consistent with the depictive theory, recent neural evidence from fMRI finds that the content of visual experience is similarly encoded across perception and imagination, since it can be decoded from the early visual cortex using multivariate cross-classification (MVCC)[33, 34]. In these studies, a classification model is first trained on the pattern of brain activity in the visual cortex elicited by viewing a set of stimuli and then tested the activity elicited by imagining the same stimuli.

Emotion regulation involves the effortful control of cognition or attention to modulate emotional reactivity. This includes both down-regulation (e.g., reducing a negative response, making it *less* negative) and up-regulation (e.g., enhancing a negative response, making it *more* negative). The effortful down-regulation of differential fear conditioning appears to produce significant down-regulation in self-reported fear[35], SCRs and activity in parts of the fear network, including the aIn[9]. Delgado et al.[9] appears to be the only previous research to evaluate the down-regulation of fear conditioning via a strategy involving imagery. During the down-regulate CS+ condition compared to the attend CS+ condition they observed reduced SCR and aIn activity along with increased activation of the dorsolateral prefrontal cortex (dlPFC). The involvement of dlPFC is consistent with broader emotion regulation findings implicating frontoparietal attention control regions in regulation[36]. There appears to be no research to date on the up-regulation of differential fear conditioning via mental imagery.

Unresolved in Delgado et al.[9] and the larger emotion regulation literature is how frontoparietal cortices produce the regulation of brain regions associated with regions of the fear network. While some research indicates that frontoparietal regions operate via connections with the ventromedial prefrontal cortex[9,37,38], other evidence indicates that regulation can occur via the modulation of perceptual areas of the occipitotemporal cortex by attention control processes initiated in frontoparietal cortices[39–42] as described by the biased-competition theory[43–45].

According to the biased-competition theory[46], when two stimuli compete for representation within a given brain region they do so in a mutually inhibitory manner. The stimulus that “wins” and becomes preferentially represented is the one that is biased either by stimulus properties or by top-down attention control processes[47]. Extending this framework to emotion regulation, when we regulate emotional stimuli using distraction or reappraisal, we are prioritizing a representation that competes with the representation of the external emotion elicitor. Combining the depictive theory with the biased-competition theory suggests, therefore, that mental imagery could facilitate emotion regulation via the activation of internally generated representations that compete with external, stimulus-driven, representations in early visual cortices. This competition would inhibit the down-stream processing of the external emotion elicitor in emotion sensitivity regions such as those of the fear network.

The present study was designed to address the following two research questions: First, can an acquired differential fear conditioned response be generated (i.e., expressed or elicited) through mental imagery, consistent with the mechanisms of the depictive theory of mental imagery? Second, can mental imagery be deployed to regulate differential fear conditioned responding via mechanisms of the depictive theory and the biased-competition model of attention? In order to address these two questions, we conducted a two-visit, two experiment, study that combined differential fear conditioning with manipulations of mental imagery. Both visits involved fMRI combined with SCR recordings and self-reported measures of fear and imagery. A retinotopic functional localizer was used to identify the regions of the early visual cortex (i.e., V1, V2, V3, V4/V3AB) and MVCC[48] was used to quantify the effects of mental imagery in the visual cortices.

The first question was addressed by testing our first two experimental hypotheses. Hypothesis 1: Imagining a CS+ versus a CS-produces a significant differential response as measured with self-report, SCR, and activation of aspects of the fear network in particular the aIn. Hypothesis 2: Consistent with the depictive theory of imagery, there will be significant decoding of imagined CS+ versus imagined CS- trials in the early visual cortex. The second question was addressed by testing experimental hypotheses 3 and 4. Hypothesis 3: Imagining the CS- while viewing the CS+ will down-regulate fear response markers (i.e., self-reported fear, SCR, and activity in fear network regions), and imagining the CS+ while viewing the CS- will up-regulate the fear response markers. Hypothesis 4: Consistent with the depictive theory and the biased competition theory, we will observe a significant reduction in decoding accuracy of the CS+ versus CS- in early visual cortex during regulation by imagery (i.e., the down-regulation versus up-regulation conditions) compared to the conditions of viewing the CS+ versus the CS-.

## RESULTS

### Experiment 1 (Visit 1) – Fear transfer to imagined stimuli

#### Self-reported fear of shock (Figure 1c), imagery vividness and imagery effort

To test the hypothesis that fear conditioning was acquired to viewed stimuli and transferred to imagined stimuli as measured using subjective markers of fear, we ran a repeated measure analysis of variance (ANOVA) on the self-reported fear of shock data with CS-Type (CS+, CS-) and Instruction (view, imagine) as within-participant variables. There was a CS-Type x Instruction interaction, *F*(1, 12) = 6.50, *p* = .025, 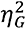 = .063, reflecting a greater difference between CS+ and CS- in the view condition, *t*(12) = 7.31, *p* < .001, *d* = 2.03, than in the imagine condition, *t*(12) = 5.50, *p* < .001, *d* = 1.53. This last effect, however, indicates a transfer of fear conditioning when imagining the CS+ versus the CS-, as measured with self-report. Additionally, a post-hoc comparison revealed significantly greater subjective fear for ‘view’ CS+ than ‘imagine’ CS+, *t*(12) = 3.18, *p* = .008, *d* = 0.88. There was also both a significant main effect of CS-Type, *F*(1, 12) = 62.91, *p* < .001, 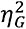 = 0.56, with higher self-reported fear for CS+ than for CS- irrespective of Instruction, and a significant main effect of Instruction, *F*(1, 12) = 7.44, *p* = .017, 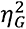 = 0.12, with greater self-reported fear for viewed than for imagined stimuli.

We verified that self-rated effort to form a mental image did not differ significantly between imagine CS+ and imagine CS- (*M* = 5.15, *SD* = 1.41, and *M* = 5.46, *SD* = 1.05, respectively), *t*(12) = 1.00, *p* = .337 (Supplemental Figure 1). Likewise, vividness of mental imagery did not significantly differ between imagine CS+ and imagine CS- (*M* = 4.31, *SD* = 1.11, range = 3-6 and *M* = 4.00, *SD* = 1.08, range = 2-6, respectively), *t*(12) = 0.88, *p* = .392 (Supplemental Figure 1). This allowed us to rule out both imagery effort and vividness as potential confounds that might explain differences in any physiological or neural differences observed when comparing ‘imagine’ CS+ to ‘imagine’ CS-. Additionally, one-sample t-tests for both CS+ imagery vividness, t(12) = 10.75, *p* < 0.001, and CS- imagery vividness, t(12) = 10.01, *p* < 0.001, confirmed that participants’ mental imagery vividness was significantly greater than ‘non-existent’ (i.e., greater than 1 on the Likert scale). Moreover, no participant selected a vividness rating less than 2, with an overall range of 2-6 across both CS+ and CS- imagery trials.

#### Electrodermal activity (Figure 1d)

To test the hypothesis that fear conditioning was acquired to viewed stimuli and transferred to imagined stimuli as measured using physiological markers of fear, we ran a similar ANOVA on the SCR. There was a significant main effect of CS-Type, *F*(1, 11) = 14.82, *p* = .003, 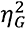 = .059, with higher SCR for CS+ than for CS- irrespective of Instruction; there was no significant main effect of Instruction, *F*(1, 11) = 2.09, *p* = .18, and no significant CS-Type x Instruction interaction, *F*(1, 11) = 0.23, *p* = .64. Although we observed no interaction and given our *a priori* interest in the differential CR within each Instruction, we conducted two follow-up t-tests for the view and imagine conditions, respectively. These tests indicated that SCRs were significantly greater for CS+ than for CS- in the view condition, *t*(11) = 3.29, *p* = .004, one-tailed, *d* = 0.95, and in the imagine condition, *t*(11) = 2.02, *p* = .034, one-tailed, *d* = 0.58 (one-tailed tests were used because we had a specific directional hypothesis regarding SCR and fear conditioning). Also, note that no significant difference was observed between CS+ view and CS+ imagine, *t*(11) = 1.16, *p* = .27, nor between CS- view and CS- imagine, *t*(11) = 1.28, *p* = .23. Together, these results suggest that fear acquired to viewed stimuli transfers to imagined stimuli, as measured with SCR.

#### Brain Imaging (fMRI)

##### Differential fear conditioning to viewed percepts (Figure 1e; Table 1)

Our fear conditioning manipulation to viewed stimuli produced canonical activation in a network of regions associated with differential fear conditioning[6]. Specifically, there was significantly greater signal in parts of bilateral anterior insula, bilateral dACC, right thalamus, right striatum, and right aspects of the midbrain during the ‘view’ CS+ compared to ‘view’ CS- condition.

##### Differential fear transfer to imagined percepts (Figure 1f)

In the critical contrast for experiment one, we observed significantly greater activity in the right aIn, right dlPFC and bilateral inferior parietal lobe during the ‘imagine’ CS+ compared to the ‘imagine’ CS- condition (Table 1).

**Table 1:**
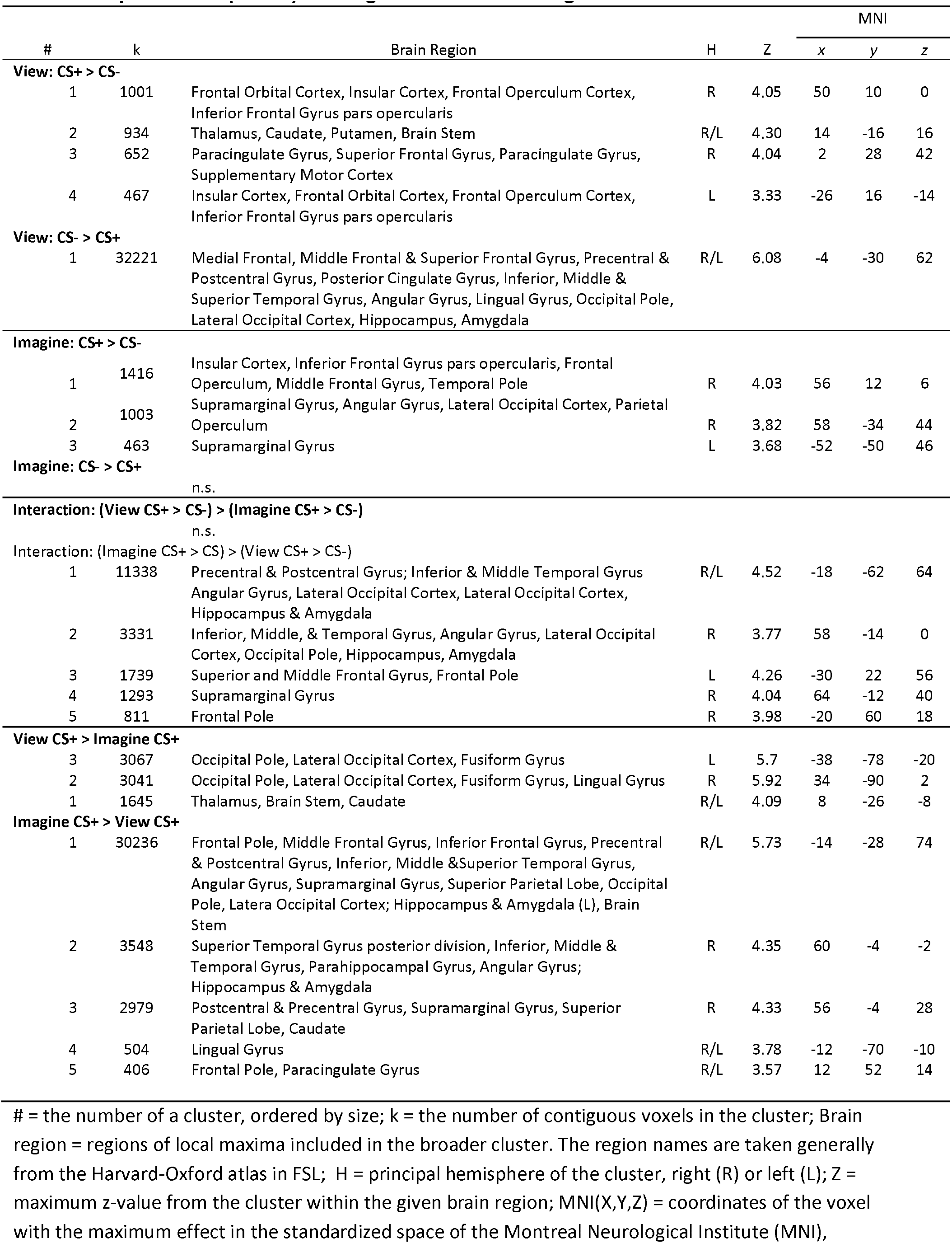
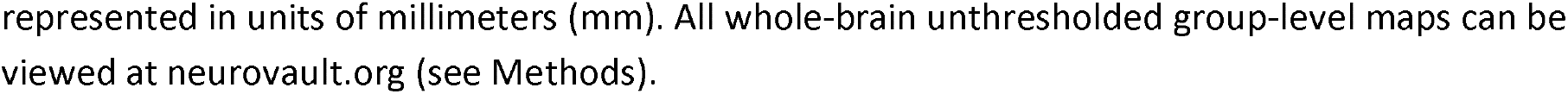
Experiment 1 (Visit 1) – Fear generalization to imagined stimuli

| # | k | Brain Region | H | Z | MNI |  |  |
| --- | --- | --- | --- | --- | --- | --- | --- |
|  |  |  |  |  | x | y | z |
| View: CS+ > CS- |  |  |  |  |  |  |  |
| 1 | 1001 | Frontal Orbital Cortex, Insular Cortex, Frontal Operculum Cortex, Inferior Frontal Gyrus pars opercularis | R | 4.05 | 50 | 10 | 0 |
| 2 | 934 | Thalamus, Caudate, Putamen, Brain Stem | R/L | 4.30 | 14 | -16 | 16 |
| 3 | 652 | Paracingulate Gyrus, Superior Frontal Gyrus, Paracingulate Gyrus, Supplementary Motor Cortex | R | 4.04 | 2 | 28 | 42 |
| 4 | 467 | Insular Cortex, Frontal Orbital Cortex, Frontal Operculum Cortex, Inferior Frontal Gyrus pars opercularis | L | 3.33 | -26 | 16 | -14 |
| View: CS- > CS+ |  |  |  |  |  |  |  |
| 1 | 32221 | Medial Frontal, Middle Frontal & Superior Frontal Gyrus, Precentral & Postcentral Gyrus, Posterior Cingulate Gyrus, Inferior, Middle & Superior Temporal Gyrus, Angular Gyrus, Lingual Gyrus, Occipital Pole, Lateral Occipital Cortex, Hippocampus, Amygdala | R/L | 6.08 | -4 | -30 | 62 |
| Imagine: CS+ > CS- |  |  |  |  |  |  |  |
| 1 | 1416 | Insular Cortex, Inferior Frontal Gyrus pars opercularis, Frontal Operculum, Middle Frontal Gyrus, Temporal Pole | R | 4.03 | 56 | 12 | 6 |
| 2 | 1003 | Supramarginal Gyrus, Angular Gyrus, Lateral Occipital Cortex, Parietal Operculum | R | 3.82 | 58 | -34 | 44 |
| 3 | 463 | Supramarginal Gyrus | L | 3.68 | -52 | -50 | 46 |
| Imagine: CS- > CS+ |  |  |  |  |  |  |  |
| n.s. |  |  |  |  |  |  |  |
| Interaction: (View CS+ > CS-) > (Imagine CS+ > CS-) |  |  |  |  |  |  |  |
| n.s. |  |  |  |  |  |  |  |
| Interaction: (Imagine CS+ > CS-) > (View CS+ > CS-) |  |  |  |  |  |  |  |
| 1 | 11338 | Precentral & Postcentral Gyrus; Inferior & Middle Temporal Gyrus Angular Gyrus, Lateral Occipital Cortex, Lateral Occipital Cortex, Hippocampus & Amygdala | R/L | 4.52 | -18 | -62 | 64 |
| 2 | 3331 | Inferior, Middle, & Temporal Gyrus, Angular Gyrus, Lateral Occipital Cortex, Occipital Pole, Hippocampus, Amygdala | R | 3.77 | 58 | -14 | 0 |
| 3 | 1739 | Superior and Middle Frontal Gyrus, Frontal Pole | L | 4.26 | -30 | 22 | 56 |
| 4 | 1293 | Supramarginal Gyrus | R | 4.04 | 64 | -12 | 40 |
| 5 | 811 | Frontal Pole | R | 3.98 | -20 | 60 | 18 |
| View CS+ > Imagine CS+ |  |  |  |  |  |  |  |
| 3 | 3067 | Occipital Pole, Lateral Occipital Cortex, Fusiform Gyrus | L | 5.7 | -38 | -78 | -20 |
| 2 | 3041 | Occipital Pole, Lateral Occipital Cortex, Fusiform Gyrus, Lingual Gyrus | R | 5.92 | 34 | -90 | 2 |
| 1 | 1645 | Thalamus, Brain Stem, Caudate | R/L | 4.09 | 8 | -26 | -8 |
| Imagine CS+ > View CS+ |  |  |  |  |  |  |  |
| 1 | 30236 | Frontal Pole, Middle Frontal Gyrus, Inferior Frontal Gyrus, Precentral & Postcentral Gyrus, Inferior, Middle & Superior Temporal Gyrus, Angular Gyrus, Supramarginal Gyrus, Superior Parietal Lobe, Occipital Pole, Lateral Occipital Cortex; Hippocampus & Amygdala (L), Brain Stem | R/L | 5.73 | -14 | -28 | 74 |
| 2 | 3548 | Superior Temporal Gyrus posterior division, Inferior, Middle & Temporal Gyrus, Parahippocampal Gyrus, Angular Gyrus; Hippocampus & Amygdala | R | 4.35 | 60 | -4 | -2 |
| 3 | 2979 | Postcentral & Precentral Gyrus, Supramarginal Gyrus, Superior Parietal Lobe, Caudate | R | 4.33 | 56 | -4 | 28 |
| 4 | 504 | Lingual Gyrus | R/L | 3.78 | -12 | -70 | -10 |
| 5 | 406 | Frontal Pole, Paracingulate Gyrus | R/L | 3.57 | 12 | 52 | 14 |

##### Interaction of differential visual fear conditioning versus imaginal fear transfer (Supplemental Figure 2; Table 1)

This interaction revealed a broad pattern of activity such that differential imaginal fear transfer [(‘imagine’ CS+) – (‘imagine’ CS-)] was significantly greater than differential visual fear conditioning [(‘view’ CS+) – (‘view’ CS-)]. This interaction effect was found in several noteworthy regions including bilateral hippocampus, amygdala, and supramarginal gyrus, though it appears driven by deactivation during view CS+ trials (see Table 1 for full details). On the other hand, no clusters were significantly greater for differential visual fear conditioning [(‘view’ CS+) – (‘view’ CS-)] compared to differential imaginal fear transfer [(‘imagine’ CS+) – (‘imagine’ CS-)].

##### Contrast of viewing versus imagining the fear conditioned percept (Supplemental Figure 3; Table 1)

We performed the contrast of ‘view’ CS+ versus ‘imagine’ CS+ to directly assess which regions are differentially recruited when a visual percept generates a fear conditioned response versus when an imagined percept generates one. As one would predict viewing the CS+ produced significantly more activity in bilateral visual cortex, thalamus, and midbrain regions of the brainstem (i.e., bottom-up regions). Conversely, imagining the CS+ produces significantly greater signal in broad parts of bilateral frontoparietal regions and bilateral medial temporal lobe areas including hippocampus and amygdala.

##### Neural overlap of differential fear conditioning and imagery transfer (Supplemental Figure 4; Table 2)

The neural overlap of the thresholded and corrected whole-brain group-level maps from the ‘view’ CS+ > ‘view’ CS- analysis and the ‘imagine’ CS+ > ‘imagine’ CS- analysis overlap the right aIn and inferior frontal gyrus. This may suggest that the right aIn is an important location for the expression of both stimulus-driven differential fear conditioning (‘view’ CS+ > ‘view’ CS-) and imagery-driven differential fear transfer (‘imagine’ CS+ > ‘imagine’ CS-).

**Table 2:** Day 1, Conditioning Phase: Neural Overlap

| Conjunction |  | H | Z | k | MNI |  |  |
| --- | --- | --- | --- | --- | --- | --- | --- |
| Brain Region |  |  |  |  | x | y | z |
| **View ( CS+ > CS-) ∩ Imagine ( CS+ > CS-)** |  |  |  |  |  |  |  |
| Anterior Insula / Frontal Operculum |  | R | 11.80 | 304 | 50 | 10 | 0 |
| Inferior Frontal Gyrus |  | R | 8.25 | 13 | 52 | 22 | 18 |

The results in this table are derived from taking the product of two independent whole-brain maps each of which was thresholded and cluster-corrected independently for multiple comparisons (see Methods for further details). H = principal hemisphere of the cluster, right (R) or left (L); Z = maximum z-value of cluster; $k$ = number of contiguous voxels in the cluster; MNI(X,Y,Z) = coordinates of the center of gravity in the standardized space of the Montreal Neurological Institute (MNI), represented in units of millimeters (mm). All whole-brain unthresholded group-level maps can be viewed at neurovault.org (see Methods).

##### PPI, whole-brain (Supplemental Figure 5, Table 3)

The PPI analysis focused on the connectivity of the right aIn during ‘view’ CS+ versus ‘imagine’ CS+. The right aIn cluster (restricted with an anatomical insula mask) was defined from the “neural overlap of differential fear conditioning and imagery transfer” (see above) and was used as the seed region for the PPI. This analysis revealed significantly stronger (and positive) functional connectivity during ‘imagine’ versus ‘view’ CS+ between right aIn and bilateral aspect of both the MTL and parietal regions, including bilateral amygdala, hippocampus, lateral occipital cortex, angular gyrus, and superior parietal lobe. On the other hand, there were no significant effects for ‘view’ CS+ stronger than ‘imagine’ CS+.

**Table 3:**
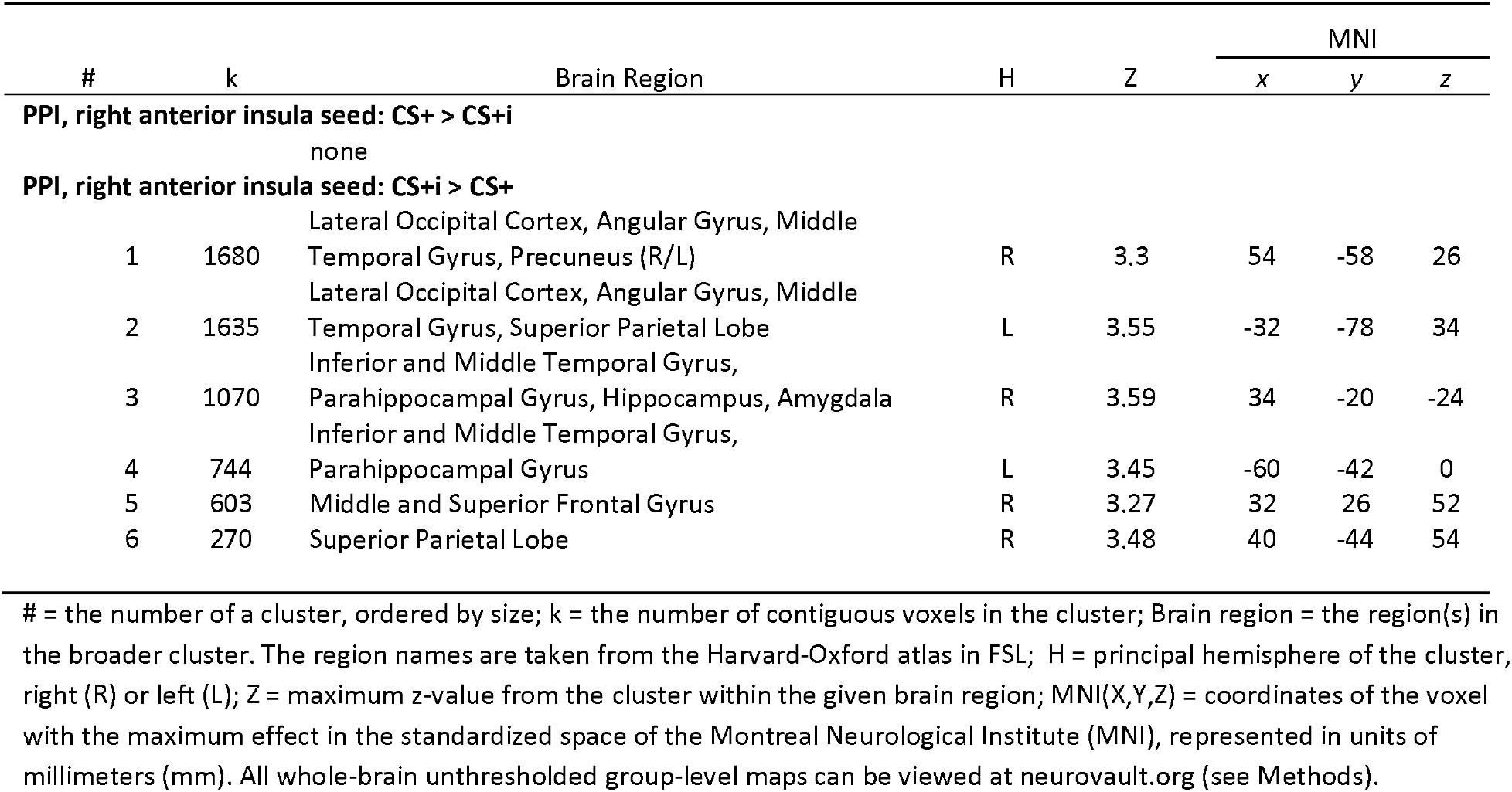
Psychophysiological interaction model with right anterior insula seed during conditioning phase

| # | k | Brain Region | H | Z | MNI |  |  |
| --- | --- | --- | --- | --- | --- | --- | --- |
|  |  |  |  |  | x | y | z |
| PPI, right anterior insula seed: CS+ > CS+i |  |  |  |  |  |  |  |
|  |  | none |  |  |  |  |  |
| PPI, right anterior insula seed: CS+i > CS+ |  |  |  |  |  |  |  |
| 1 | 1680 | Lateral Occipital Cortex, Angular Gyrus, Middle Temporal Gyrus, Precuneus (R/L) | R | 3.3 | 54 | -58 | 26 |
| 2 | 1635 | Lateral Occipital Cortex, Angular Gyrus, Middle Temporal Gyrus, Superior Parietal Lobe | L | 3.55 | -32 | -78 | 34 |
| 3 | 1070 | Inferior and Middle Temporal Gyrus, Parahippocampal Gyrus, Hippocampus, Amygdala | R | 3.59 | 34 | -20 | -24 |
| 4 | 744 | Inferior and Middle Temporal Gyrus, Parahippocampal Gyrus | L | 3.45 | -60 | -42 | 0 |
| 5 | 603 | Middle and Superior Frontal Gyrus | R | 3.27 | 32 | 26 | 52 |
| 6 | 270 | Superior Parietal Lobe | R | 3.48 | 40 | -44 | 54 |

##### Multivariate Cross-Classification in early visual cortex (Figure 2)

To quantify and validate that the participants followed the imagery instructions when cued to do so, we measured the cross-classification accuracy of models trained to discriminate the CS+ from the CS- using data from the classifier training phase and tested the data from the conditioning phase (Figure 2a) for each of our visual cortex ROIs. Not surprisingly, for the ‘view’ conditions our classification accuracy was significantly above chance for all regions, V1 (acc = 59.38%; *p* = .0017), V2 (acc = 67.36%; *p* < .0001), V3 (acc = 66.67%; *p* < .0001), and V4-V3AB (acc = 61.81%; *p* < .0001). Importantly, we also found significant classification during the ‘imagine’ conditions in V2 (acc = 56.60%; *p* = .0082), V3 (acc = 60.07%; *p* = .0002), and V4-V3B (acc = 59.38%; *p* = .0003). We also found that classifier accuracy was significantly better for the ‘view’ conditions compared to the ‘imagine’ conditions in V1 (*p =* .0166), V2 (*p =* .0048), and V3 (*p =* .0495), though there was no significant difference found in V4-V3AB (*p =* .2812). Overall, these findings provide group-level evidence that participants were able to generate a pattern of activation during mental imagery similar to a pattern of activation generated when viewing the CS+ versus CS- and are consistent with the depictive theory.

### Experiment 2 (Visit 2) – Regulation of fear conditioning via mental imagery

#### Self-reported fear of shock (Figure 3c), imagery vividness and imagery effort

To test the hypothesis that mental imagery can be used to regulate a fear conditioned response as measured using a subjective marker of fear, we ran a repeated-measures ANOVA on the self-reported fear of shock with CS-Type (CS+, CS-) and Instruction (‘view’, ‘regulate’) as within-participant variables. Critically, this revealed a CS-Type x Instruction interaction, *F*(1, 11) = 72.40, *p* < .001, 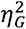 = .617, reflecting significantly more subjective fear for the CS+ versus CS- in the view condition, *t*(11) = 8.51, *p* < .001, *d* = 2.46, and the reverse effect in the regulate condition such that participants reported more fear for the ‘up-regulate’ (vCS-/iCS+) versus ‘down-regulate’ (vCS+/iCS-) condition, *t*(11) = 3.22, *p* = .008, *d* = 0.93. Additionally, whereas there was a significant reduction in subjective fear during ‘down-regulate’ compared to ‘view’ CS+, *t*(11) = -9.916, *p* = .008, *d* = 2.86, there was a significant increase in fear during ‘up- regulate’ versus ‘view’ CS-, *t*(11) = 2.245, *p* = .032, *d* = 0.71. There was also a significant main effect of CS-Type, *F*(1, 11) = 29.24, *p* < .001, 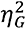 = .357, with higher self-reported fear for CS+ than for CS-, and a significant main effect of Instruction, *F*(1, 11) = 33.00, *p* < .001, 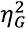 = .381, with greater self-reported fear for viewed than for imagined stimuli. These results suggest that mental imagery is capable of leading to both the down-regulation of a visual fear conditioned stimulus, and the up-regulation of fear by imagining a CS+ as measured by self-report.

We verified that effort to form a mental image did not differ significantly between imagining the CS+ (i.e., ‘up-regulate’) and imagining the CS- (i.e., ‘down-regulate) during the regulation phase (*M* = 6.08, *SD* = 0.90, and *M* = 5.75, *SD* = 1.06, respectively), *t*(11) = 1.48, *p* = .166 (Supplemental Figure 6 – Left). Likewise, vividness of mental imagery was not significantly different when imagining the CS+ versus imagining the CS- (*M* = 4.83, *SD* = 1.27, range = 3-7, and *M* = 4.58, *SD* = 1.31, range = 3-7, respectively), *t*(11) = 1.15, *p* = .275 (Supplemental Figure 6). Additionally, one-sample t-tests for both CS+ imagery vividness, t(11) = 10.48, *p* < 0.001, and CS- imagery vividness, t(11) = 9.47, *p* < 0.001, confirmed that participants’ mental imagery vividness was significantly greater than ‘non-existent’ (i.e., greater than 1 on the Likert scale). Moreover, imagery vividness across both imagery conditions was never less than 3 on the 7-point Likert scale.

#### Electrodermal activity (Figure 3d)

To test the hypothesis that mental imagery can be used to regulate a fear conditioned response as measured using a physiological marker of fear, similar to the self-report analysis we ran an ANOVA on the SCR data. This revealed a CS Type x Instruction interaction, *F*(1, 8) = 5.70, *p* = .044, 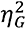 = .038. Subsequent t-tests indicated that the SCR was significantly greater for CS+ than for CS- in the view condition, *t*(8) = 3.01, *p* = .017, *d* = 1.00, but no significant difference between CS+ and CS- was observed in the regulate condition, *t*(8) = 0.22, *p* = .42. In addition, compared to the ‘view’ CS+ condition, the ‘regulate’ CS+ condition (i.e., vCS+/iCS-) had a significantly lower SCR, *t*(8) = 3.38, *p* = .01, *d* = 0.86. The ANOVA also revealed both a main effect of CS-Type, *F*(1, 8) = 5.97, *p* = .04, 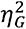 = .047, with greater SCR for CS+ than for CS-, and a main effect of Instruction, *F*(1, 8) = 6.34, *p* = .04, 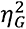 = .079, with greater SCR for the ‘view’ conditions compared to the ‘regulate’ conditions. Overall, this provides evidence for the use of mental imagery in the down-regulation of a fear conditioned response but no evidence for the up-regulation of fear as measured with SCR.

#### Brain imaging (fMRI)

##### Univariate region-of-interest analysis (right aIn; Figure 3e,f)

Using the right aIn mask produced from the “neural overlap of differential fear conditioning and imagery transfer” in Experiment 1, we found that fear-associated reactivity is modulated by mental imagery-based emotion regulation. Specifically, we ran a 2(CS-Type: CS+, CS-) by 2 (Instruction: ‘view’, ‘regulate’) repeated measures ANOVA on mean percent signal change extracted from the right aIn. This revealed the key interaction, *F*(1, 11) = 22.48, *p* < .001, 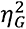 = .282. This interaction was driven by a significant down-regulation of activity from ‘view’ CS+ to ‘regulate’ CS+ (vCS+/iCS-), *t*(11) = 2.64, *p* = .023, *d* = 0.761, and a significant up-regulation of activity from ‘view’ CS- to ‘regulate’ CS- (vCS-/iCS+), *t*(11) = 3.46, *p* = .005, *d* = 0.999. Furthermore, there was significantly more activation of right aIn when viewing the CS+ compared to viewing the CS-, *t*(11) = 3.95, *p* = .002, *d* = 1.104, and there was more activity when regulating the CS- compared to regulating the CS+, *t*(11) = 2.961, *p* = .013, *d* = 0.855.

##### Differential fear conditioning to viewed percepts (Figure S7; Table 4)

As with Experiment 1, a whole-brain analysis with Experiment 2 revealed significantly greater activation for ‘view’ CS+ compared to ‘view’ CS- in a network of regions associated with fear conditioning[6] including bilateral aIn, dACC, thalamus, and midbrain regions of the brainstem. Conversely, we also found several areas associated with more activity for ‘view’ CS- versus ‘view’ CS+ such as left MTL.

##### Contrast of the ‘down-regulation’ versus ‘up-regulation’ of a fear conditioned response

The whole-brain analysis directly comparing the two ‘regulate’ conditions revealed no significant differences between the ‘down-regulation’ (viewing the CS+ while imagining the CS-) versus ‘up-regulation’ of a fear conditioning (viewing the CS- while imagining the CS+).

##### Contrast of ‘down-regulation’ versus ‘view’ CS+ (Figure 4a; Table 4)

The first of two key univariate whole-brain contrasts of the regulation phase compared viewing the CS+ to down-regulation of the CS+ (vCS+/iCS-). This revealed a robust reduction in brain activity across large parts of the fear conditioning network, including the bilateral anterior insula, bilateral ventral striatum, bilateral thalamus, and midbrain aspects of the brainstem putatively including the PAG during down-regulation compared to viewing the CS+. On the other hand, down-regulation compared to viewing the CS+ was associated with greater activation in posterior aspects of the frontoparietal attention networks, specifically bilateral superior parietal lobe. There was also greater activity in aspects of bilateral medial postcentral gyrus and left lateral occipital cortex.

##### Contrast of ‘up-regulation’ versus ‘view’ CS- (Figure 4b; Table 4)

The second key univariate whole-brain contrast of the regulation phase compared viewing the CS- to the CS- up-regulation condition (vCS-/iCS+). This revealed an increase in brain activity across some parts of the fear conditioning network, such as the bilateral aIn and bilateral striatum. Additionally, there was greater activation in aspects of frontoparietal attention areas, including bilateral DLPFC, bilateral IFG, and bilateral superior parietal lobe. Conversely, viewing the CS- compared to the up-regulation condition (vCS-/iCS+) was associated with greater activity is regions associated with the default-mode network, such as bilateral ventromedial prefrontal cortex, and bilateral posterior cingulate cortex.

Interaction of Stimulus Type (CS+, CS-) by Instruction (View, Regulate), (Supplemental Figure 8; Table 4)

**Table 4:**
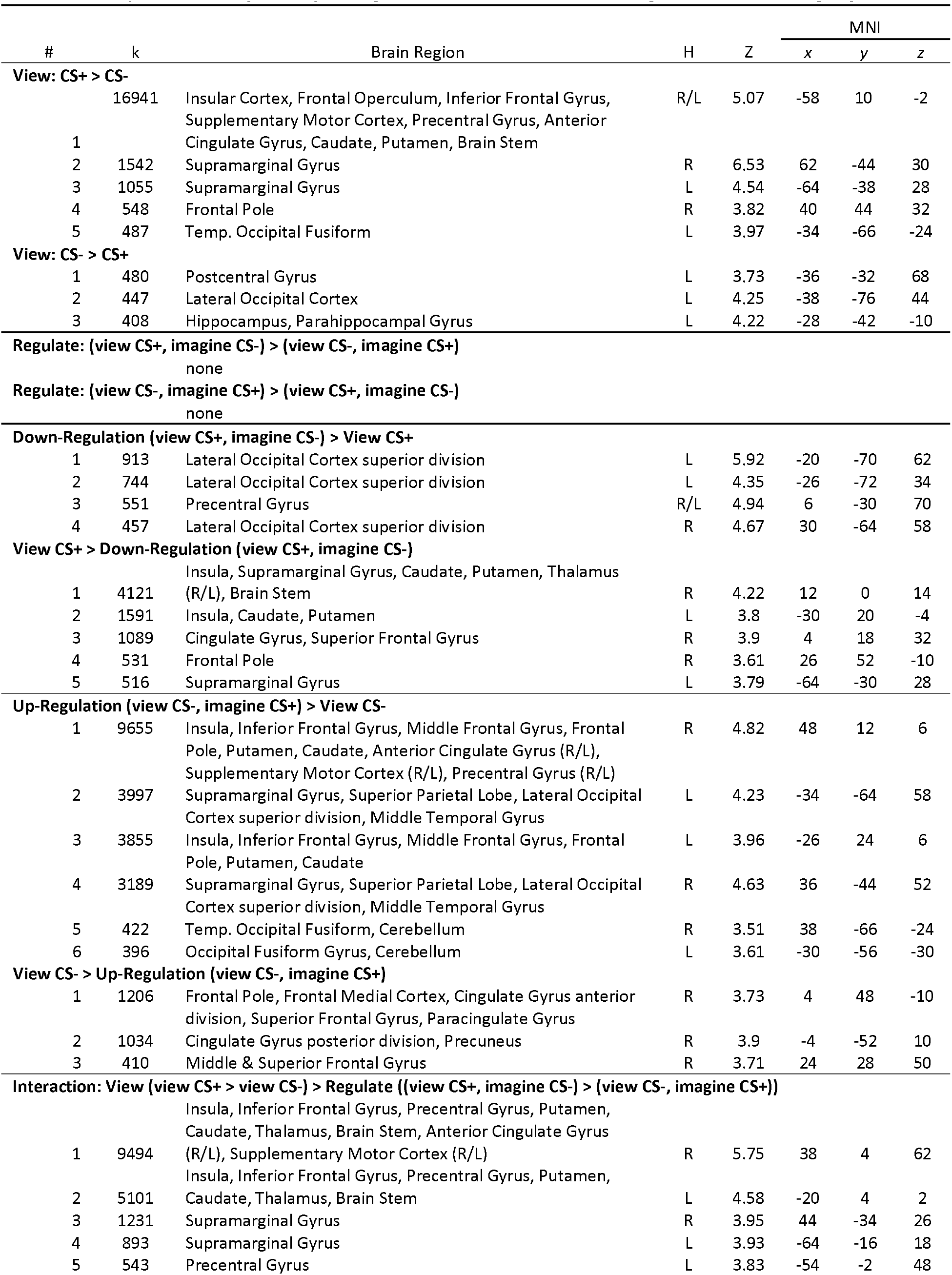
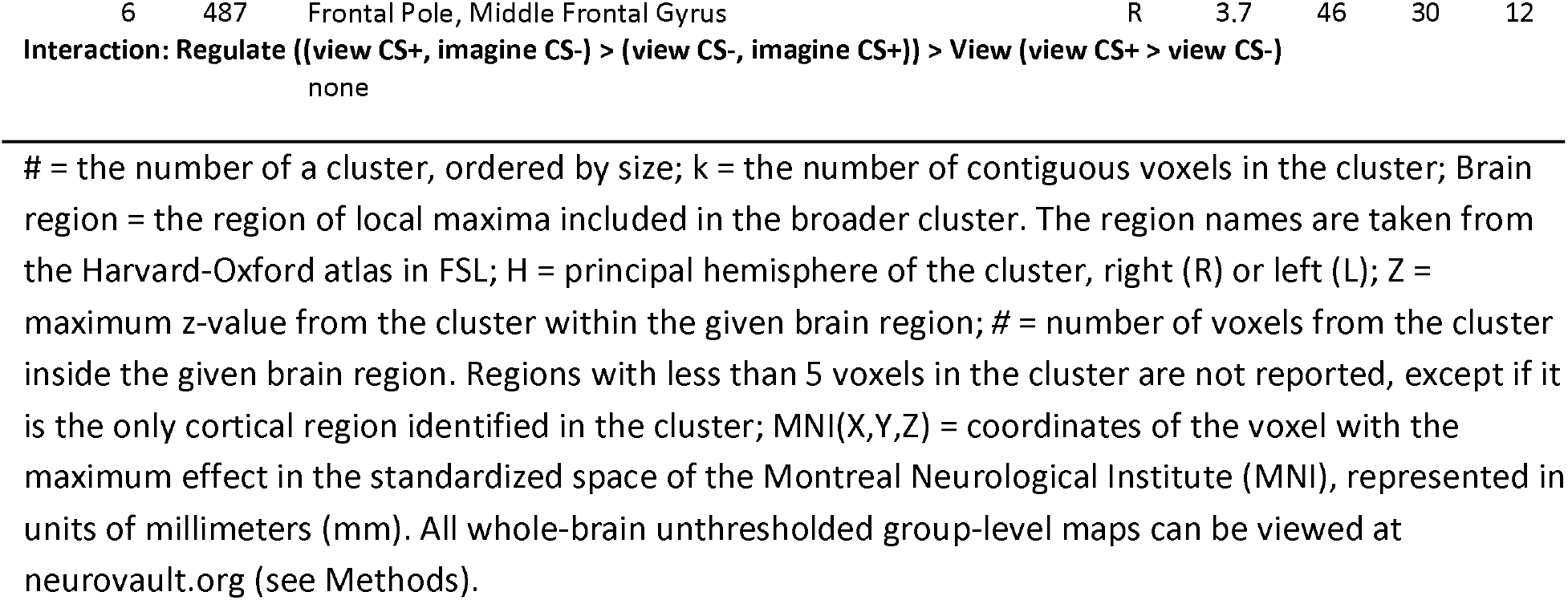
Experiment 2 (Visit 2) – Regulation of fear conditioning via mental imagery

As predicted this interaction revealed significant modulation of several core fear network regions by imagery-based regulation. The general pattern of activation observed in all regions was a cross-over interaction such that activation was reduced from ‘view’ CS+ to ‘down-regulate’ CS+ and enhanced from ‘view’ CS- to ‘up-regulate’ CS-. This pattern was observed in bilateral aIn, ventral striatum, and thalamus, and in midbrain aspects of the brainstem including putative parts of the periaqueductal gray (PAG).

##### PPI, whole-brain (Supplemental Figure 9; Table 5)

On the data from the regulation phase we carried out two PPI analyses using the right aIn ROI as the seed, which was from the differential fear conditioning neural overlap analysis [(vCS+ > vCS-) ∩ (iCS+ > iCS-)] performed on the conditioning phase (Visit 1) data. The first PPI analysis compared aIn connectivity during ‘down-regulate’ CS+ compared to ‘view’ CS+. This revealed significantly greater functional connectivity between right aIn and right amygdala, insula and IFG during ‘down-regulate’ CS+ compared to ‘view’ CS+. There were no significant effects for ‘view’ CS+ > ‘down-regulate’ CS+. The second PPI analysis compared aIn connectivity during ‘up-regulate’ CS- compared to ‘view’ CS-. This revealed significantly greater functional connectivity between right aIn and bilateral posterior cingulate cortex, precuneus, and medial occipital cortex. There were no significant effects for ‘up-regulate’ CS- > ‘view’ CS-.

**Table 5:** Psychophysiological interaction model with right anterior insula seed during regulation phase

| # | k | Brain Region | H | Z | MNI |  |  |
| --- | --- | --- | --- | --- | --- | --- | --- |
|  |  |  |  |  | x | y | z |
| Down-Regulate (vCS+/iCS-) > View CS+ |  |  |  |  |  |  |  |
| 1 | 663 | Amygdala, Putamen, Insula | R | 3.41 | 14 | 2 | -14 |
| 2 | 395 | Angular Gyrus, Middle Temporal Gyrus | L | 3.27 | -60 | -42 | 0 |
| 3 | 355 | Lateral Occipital Cortex superior division | R | 3.15 | 10 | -66 | 64 |
| 4 | 350 | Frontal Pole, Inferior Frontal Gyrus | R | 3.37 | 38 | 40 | -8 |
| View CS+ > Down-Regulate (vCS+/iCS-) |  |  |  |  |  |  |  |
|  |  | none |  |  |  |  |  |
| Up-Regulate (vCS-/iCS+) > View CS- |  |  |  |  |  |  |  |
|  |  | none |  |  |  |  |  |
| View CS- > Up-Regulate (vCS-/iCS+) |  |  |  |  |  |  |  |
| 1 | 534 | Precuneus, Posterior Cingulate Gyrus | R/L | 3.43 | -2 | -54 | 30 |
| 2 | 388 | Supracalcarine Cortex, Intracalcarine Cortex | R/L | 3.29 | 8 | -84 | 4 |

of contiguous voxels in the cluster; Brain region = the region of local maxima included in the broader cluster. The region names are taken from the Harvard-Oxford atlas in FSL; H = principal hemisphere of the cluster, right (R) or left (L); Z = maximum z-value from the cluster within the given brain region; MNI(X,Y,Z) = coordinates of the voxel with the maximum effect in the standardized space of the Montreal Neurological Institute (MNI), represented in units of millimeters (mm). All whole-brain unthresholded group-level maps can be viewed at neurovault.org (see Methods).

##### MVCC in early visual vortex (Figure 5)

After Experiment 1 (Conditioning phase) revealed that we could decode the contents of participants’ mental imagery, in Experiment 2 we tested the prediction that imagining a stimulus competes with the representation of a stimulus being viewed such that decoding accuracy is reduced. For the ‘view’ conditions of Experiment 2 our classification accuracy was significantly above chance for V1 (acc = 61.88%; *p* < .0001), V2 (acc = 69.06%; *p* < .0001), and V3 (acc = 67.03%; *p* < .0001), though it was not significantly above chance in V4-V3AB (acc = 53.13%; *p* = .0841). During the ‘regulation’ conditions, in which one of the two Gabor stimuli is on the screen while participants imagined the opposite stimulus, we also found above chance classification accuracy (of the stimulus being viewed) for V1 (acc = 61.88%; *p* < .0001), V2 (acc = 58.49%; *p* = .0008), and V3 (acc = 60.42%; *p* < .0001), though not V4-V3AB (acc = 54.27%; *p* = .0528). Importantly, while we observed no significant differences in classification accuracy in V1 (*p* = .5069), we observed significantly reduced classification accuracy when regulating compared to viewing in V2 (*p =* .0021) and V3 (*p* = .0368). Overall, these findings indicate that regulation via mental imagery can affect classification performance within V2 and V3, presumably by disrupting the representation of the viewed stimulus.

##### MVCC in the amygdala (Figure 6)

Bach et al.[49] argued that the CS-US associations, and therefore the differential encoding of the CS+ versus the CS-, are sparsely distributed throughout the amygdala, which they demonstrated using multivariate pattern analysis. Although we observed neither evidence of greater amygdala activity for the CS+ relative to the CS- conditions (Experiment 1), nor modulation of amygdala activity during the regulation of fear conditioning (Visit 2) according to our univariate whole-brain analysis, we applied MVCC to test the prediction that mental imagery modulates the representation of CS+ versus CS-[49]. As a positive control, we first demonstrated that a model trained on ‘view’ CS+ versus ‘view’ CS- trials from the conditioning phase (Visit 1) could significantly classify ‘view’ CS+ versus ‘view’ CS- trials in the regulation phase (Visit 2) above chance, *p =* .0095. Importantly, when this model was tested on ‘down-regulate’ CS+ versus ‘up-regulate’ CS- the classification was not significantly different than chance, *p* = .7349. We also observed significantly greater classification accuracy for the ‘view’ trials compared to the ‘regulate’ trials (*p* = .0169). Overall, these findings indicate the mental imagery can disrupt the differential patterns of activity associated with the CS+ and the CS-.

## DISCUSSION

The current study evaluated the general prediction that mental imagery can be used to generate (or express) and regulate emotional reactivity. Specifically, emotional reactivity was operationalized as fear conditioned responses. The study entailed a two-visit, two experiment, study that combined differential fear conditioning with mental imagery of the conditioned stimuli. Both visits involved fMRI combined with SCR recordings and self-reported measures of fear and imagery. The effects of mental imagery were quantified using MVCC on data from regions of the early visual cortex, which were defined using a retinotopic functional localizer. Specifically, two research questions were considered and four study specific hypotheses, two related to the first research question and two related to the second research question.

### Generation of a fear conditioned response via mental imagery

Regarding the first research question, our results from Experiment 1 (Visit 1) indicate that a differential fear conditioned response is generated through mental imagery, consistent with the mechanisms of the depictive theory of mental imagery and previous research[26]. Consistent with hypothesis 1 we observed mental imagery of the CS+ versus the CS- produced a significant differential response in self-reported fear of shock, SCR, and activation of the right aIn. This pattern of effects was similar to the pattern observed in our control condition involving the visual presentation of the CS+ and CS- (without reinforcement) with several noteworthy observations. We observed that self-reported fear was significantly greater when viewing compared to imagining the CS+, which indicates that participants experience more fear towards the actual object that was paired with shock. Interestingly, this pattern was not observed with the SCR data in which case there was no interaction effect between the viewed versus imagined stimuli. Instead, we observed a main effect of CS-type such that the CS+ was associated with greater SCR than the CS- irrespective of whether participants were viewing or imagining the stimuli. Additionally, follow-up t-tests motivated from our *a priori* interests and predictions revealed significant differential SCR for the CS+ versus CS- trials when both viewing and imagine the stimuli, respectively. Future research is required to evaluate whether this lack of a differences is reliable or due to our small sample and the noisy nature of SCR. In the fMRI data, while viewing the CS+ versus CS- produced activation of bilateral aIn and dACC, imagery of the CS+ compared to CS- produced significant activation of the right aIn and more dorsal and posterior aspects of the fear network from Fullana et al.[6] including bilateral supramarginal gyri in the parietal lobe.

While no studies have evaluated the ability of imagery to generate differential fear conditioning along with the potential underlying neural mechanisms, our findings are consistent with several other areas of research relating to conditioning. For example, recent psychophysiological and fMRI research has demonstrated the potential for imagery of the CS+ to contribute to fear extinction[27,29,50] or fear reconsolidation[28, 30]. Together with the findings from Experiment 1 of the present study, these previous findings are consistent with the ability of mental imagery to contribute to the expression of differential fear conditioning. On the other hand, it is unclear from the present study how mental imagery interacts with the acquisition (i.e., the learning mechanisms) of differential conditioning. For example, Lewis et al.[26] used evaluative conditioning to demonstrate that conditioning to imagined CSs generalizes to instances of viewing those same stimuli using only reaction time measures. Additionally, two recent behavioural studies found that imagery of an aversive US can contribute to measures of associative learning[51, 52]. Future research is required to evaluate how imagery interacts with other elements of fear conditioning, such as the learning processes itself, including the neural correlates of such interactions.

The amygdala was not more active during either view or imagery CS+ trials compared to their respective CS- trials in Experiment 1. Several factors might account for our lack of sustained differential amygdala activity. The amygdala may be more strongly involved in the acquisition rather than the expression or regulation of differential conditioning[21,53–55], and as such more amygdala activity is observed early but not late in acquisition[18, 56] and when the predicted outcome of CSs are unreliable versus reliable[57]. In our study, we were primarily concerned with the expression (and later the regulation) of differential fear conditioning rather than the mechanisms of differential conditioning per se. We maintained a consistent reinforcement rate of 50% for CS+ view trials throughout both experiments (Day 1 and Day 2) and we were able to use the view conditions during both days as positive controls demonstrating the persistence fear conditioning in terms of self-report, SCR and differential brain activity in parts of the fear network. Nevertheless, the lack of heightened amygdala activity (CS+ > CS-) in the present study is also consistent with two recent meta-analyses of fMRI studies relating to differential fear conditioning[6] and differential instructed fear conditioning [58], neither of which found reliable amygdala activity. Moreover, our findings of differential activity in the aIn during both the view and imagery conditions are consistent with previous research regarding the expression of fear conditioning and the role of aIn in representing the bodily sensations associated with threat perception[23].

On the other hand, the univariate PPI analysis found significantly greater functional connectivity between the aIn and the amygdala when imagining the CS+ compared to viewing the CS+. Tracer research in non-human primates indicates that there are direct reciprocal connections between the aIn and the amygdala[59]. In humans, greater aIn-amygdala functional connectivity appears associated with individual differences in state anxiety[60], and appears elevated during fear conditioning in those with genetic vulnerability for disorders of emotion[61]. The present findings of greater aIn-amygdala connectivity might be due to the more active, top-down, processes required to imagine the CS+, which are similar to the anxious anticipation of potential threat, compared to the more passive, bottom-up processes associated with viewing the CS+. Alternatively, recent research also indicates that the aIn and the amygdala are both active during threat-related uncertainty[62]. Our finding might reflect the greater uncertainty of outcomes that was associated with imagining the CS+ compared to viewing the CS+, as participants were never told explicitly which trials or conditions would involve the US, though the US only occurred when viewing the CS+. This PPI analysis more generally revealed greater connectivity between the aIn and bilateral aspects of the MTL and parietal lobe. This more general pattern of connectivity could reflect the greater memory and attention selection processes required of mental imagery compared to passive viewing[31].

Using MVCC to test our second hypothesis we found significant decoding during mental imagery trials in V2, V3 and V4-V3AB, though in V2 and V3 the decoding accuracy was greater for viewed than imagined stimuli, which is consistent with previous research[33, 63]. This result indicates that participants followed the instruction to imagine the conditioned stimuli when cued to do so. Furthermore, these general decoding effects of visual mental imagery along with the significantly more accurate decoding of visually presented stimuli are consistent with the depictive theory of mental imagery, which predicts that mental imagery is a weaker mode of sensory perception compared to externally viewing the same object[31]. Also consistent with the depictive theory, our PPI analysis revealed significantly greater functional connectivity between the right anterior insula and bilateral MTL and superior parietal lobes when imagining the CS+ compared to viewing the CS+, as has been observed previously[31,64,65]. Participants also self-reported having a subjective experience of ‘seeing in the mind’s eye’ in that their imagery vividness for both the CS+ and CS- was significantly higher than “Non-existent”, ranging from 2-7 on the 7-point Likert scale.

One effect we did not observed was differential univariate activity in the early visual cortices when either viewing or imagining the conditioned stimuli. While some previous research has found greater activity in the early sensory regions of the conditioned stimuli for the CS+ compared to CS-[66, 67], such effects were not observed in a recent meta-analysis of human fMRI fear conditioning studies[6]. These discrepant findings could be due to differences in the types of conditioned stimuli used throughout these studies, or due to other factors such as attention or awareness of the conditioned stimuli[66,68,69].

### Regulation of a fear conditioned response with mental imagery

Regarding the second research question, our results from Experiment 2 (Visit 2) demonstrated that mental imagery can be deployed to regulate differential fear conditioned responding via mechanisms of the depictive theory and the biased-competition model of attention. In support of Hypothesis 3, imagery of the CS- while viewing the CS+ (i.e., the down-regulate condition) compared with viewing the CS+ significantly reduced self-reported fear, SCR, and activity in the right aIn ROI as well as other aspects of the fear network including bilateral anterior insula/frontal operculum, ventral striatum, thalamus, and midbrain. Moreover, in partial support of Hypothesis 3, imagery of the CS+ while viewing the CS- (i.e., the up-regulate condition) compared to viewing the CS- significantly increased self-reported fear and activity in the right aIn ROI. Imagining the CS+ increased activity in other fear network regions including bilateral anterior insula/frontal operculum, striatum, and thalamus. The only inconsistency with Hypothesis 3 was the lack of SCR up-regulation by imagery of the CS+ while viewing the CS-. The pattern of down-regulation observed in the present study is consistent with similar research on the down-regulation of differential fear conditioning using an imagery-based strategy[9]. Regarding up-regulation, to our knowledge our study is the first to measure the up-regulation of differential conditioning via imagery of the CS+. However, previous research has found that reappraisal can up-regulate one’s emotional response to fear conditioned stimuli as measured both by self-report and increased heart rate and pupil dilation[12]. In addition, while we found no evidence for the up-regulation of the SCR, previous research has found that the up-regulation of negative affect while viewing threat-related images is associated with an increased SCR[70].

Frontoparietal regions are generally associated with the cognitive control of emotion. Consistent with previous findings our study revealed that aspects of the superior parietal lobe were more active during both down- and up-regulation compared to their respective viewing conditions. This observation is consistent with the role of superior aspects of the parietal lobe being involved in aspects of visual mental imagery[64, 71] and emotion regulation via reappraisal and distraction[72, 73].

As in Experiment 1, there was no univariate increase in amygdala activation when viewing the CS+ compared to the CS- nor was there modulation of the amygdala via imagery during the regulation conditions. Consistent with previous research[49, 74], using MVCC, a model trained to distinguish CS+ from CS- using the pattern of bilateral amygdala activity during Experiment 1 (viewing CS+ vs. view CS-) was able to significantly decode the presence of the visual CS+ versus CS- during Experiment 2. Importantly, this same model was not able to decode the CS+ from the CS- during the regulation conditions despite identical visual stimuli. Overall, these findings indicated that mental imagery can successfully modulate differential fear conditioning, which is consistent with the results obtained in our measures of fear.

Regarding the fourth and final hypothesis: consistent with both the depictive theory and the biased competition theory, we observed a significant reduction in decoding accuracy of the CS+ versus CS- in V2 and V3 during regulation by imagery (i.e., the down-regulation versus up- regulation conditions) compared to the conditions of viewing the CS+ versus the CS-. These effects were observed along with the commensurate modulation of our three measures of fear as noted above. Combined with the findings from Experiment 1 in which we observed significant decoding in early visual cortices during imagery, our findings from Experiment 2 are consistent with previous research suggesting that the regulation of emotional reactivity can occur via frontoparietal attention connections to early perceptual and sensory areas, such as the occipitotemporal areas involved in vision[5,39,44,75]. There is, however, debate in the literature regarding how early and in which regions attentional competition associated with the regulation of emotion takes place. While some suggest that it occurs in multimodal areas of the temporal lobe reflecting a late process [5, 44], other have suggested that the effects of competition can also occur in early sensory regions such as early visual cortex[39,43,75,76]. Our results indicate that mental imagery can impact neural representations in the early visual cortices, as early as V2 and V3. This does not imply that competition for the purposes of emotion regulation cannot occur later in information processing, but future research is required to determine how different factors (e.g., using stimuli such as objects or words) affect where and when competition occurs. Finally, while the emotion regulation manipulation used in the present study is best considered a form of distraction, some forms of cognitive reappraisal rely on facets of mental imagery[3]. For example, when reappraisal involves imagining novel aspects to a scene it may operate via mechanisms of mental imagery and biased-competition along the occipitotemporal pathways.

The present findings do not refute the likely presences of anterior regulatory pathways involving the lateral PFC [9,37,38] and its connections to the amygdala and aIn. Rather, they are indicative of additional and complementary regulatory mechanisms involving attention selection [43, 80]. An additional possibility is that during down-regulation (vCS+/iCS-), imagery of the CS- is acting as a conditioned inhibitor as in a summation test. In rodent research such conditioned inhibition involves the depression of amygdala activity [77], which could explain the MVCC results we observed in the amygdala. However, this account cannot explain why the decoding accuracy in V2 and V3 during regulation by imagery was significantly lower than the decoding accuracy of the view conditions. Future research in both human and non-human animals might consider the role of sensory cortices and their modulation by conditioned inhibitors as has been done with excitatory conditioned stimuli [78, 79].

### Other considerations

The relatively small sample size might be considered a limitation of the current study. However, across both Experiment 1 and 2 we observed consistent and robust differential conditioned responses such that ‘view’ CS+ was greater than ‘view’ CS- for all dependent measures emphasized in the analyses of mental imagery. From Experiments 1 and 2 this included the measures of self-reported fear (Figures 1c & 3c), SCR (Figures 1d & 3d), univariate BOLD response in the fear network including bilateral aIn and dACC (Figures 1e & 3e, and Supplemental Figure 7), and the MVCC decoding in the early visual cortex (V1-V3, Figures 2b & 5b). This also included the amygdala decoding (Figure 6b). Taken together, the robust positive control effects alongside the multiple convergent results from across the dependent measures suggest that the primary results regarding the impact of mental imagery on the generation and regulation of emotion via the depictive and biased competition theories are valid.

**Figure 1.**
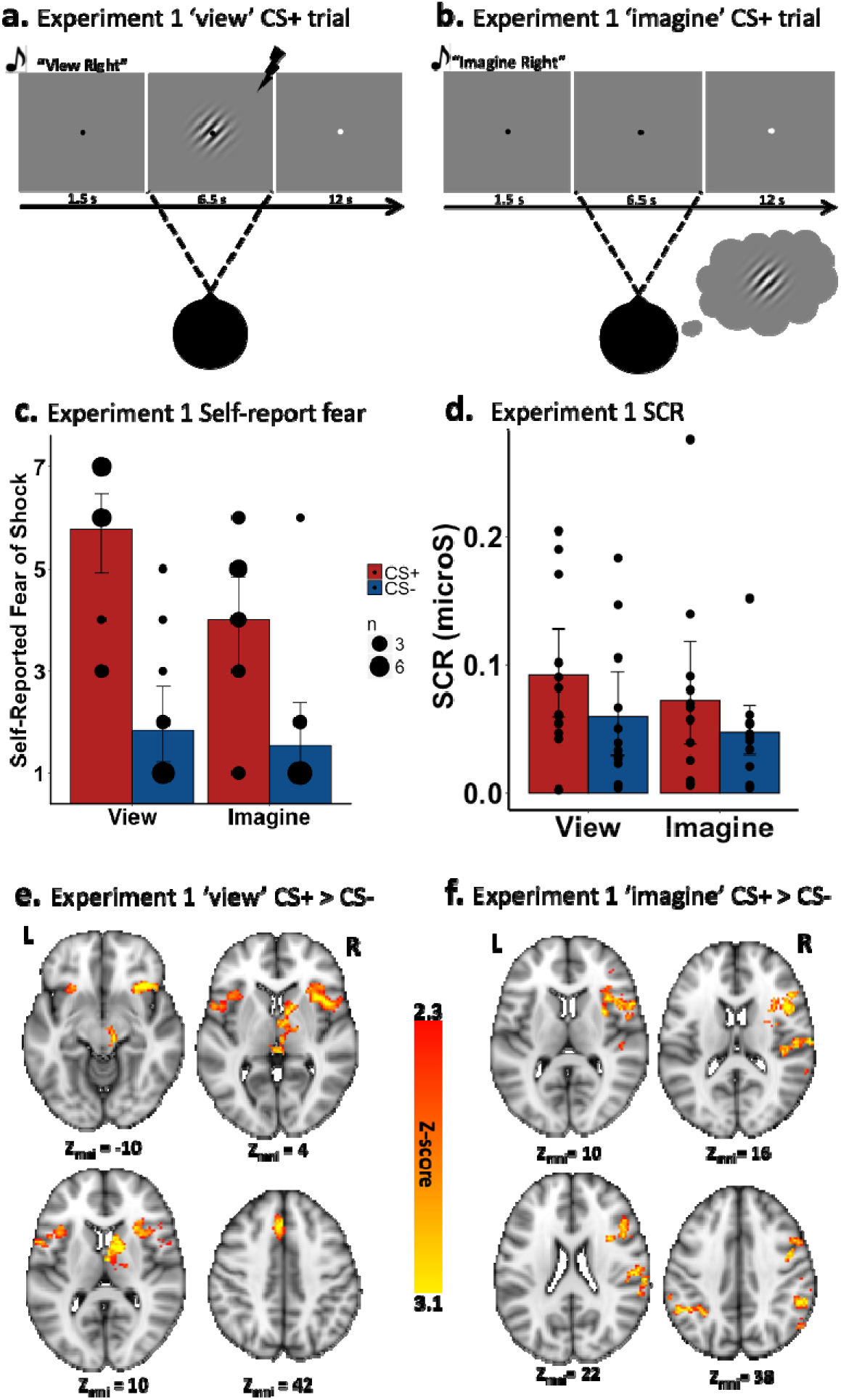
The trial sequence from Experiment 1 for (a) a ‘view’ CS+ trial and (b) an ‘imagine’ CS+ trial. c) Self-reported fear of shock; and d) SCR results. While Red bars denoted the CS+ and Blue bars represented the CS-. Error-bars represent 95% confidence intervals. Black dots represent individual data points. For the self-reported fear of shock data, the size of the circle represents the number of participants that endorsed a given response. Activation maps for (e) ‘view’ CS+ > ‘view’ CS-. Activation maps for (f) ‘imagine’ CS+ > ‘imagine’ CS-.

**Figure 2.**
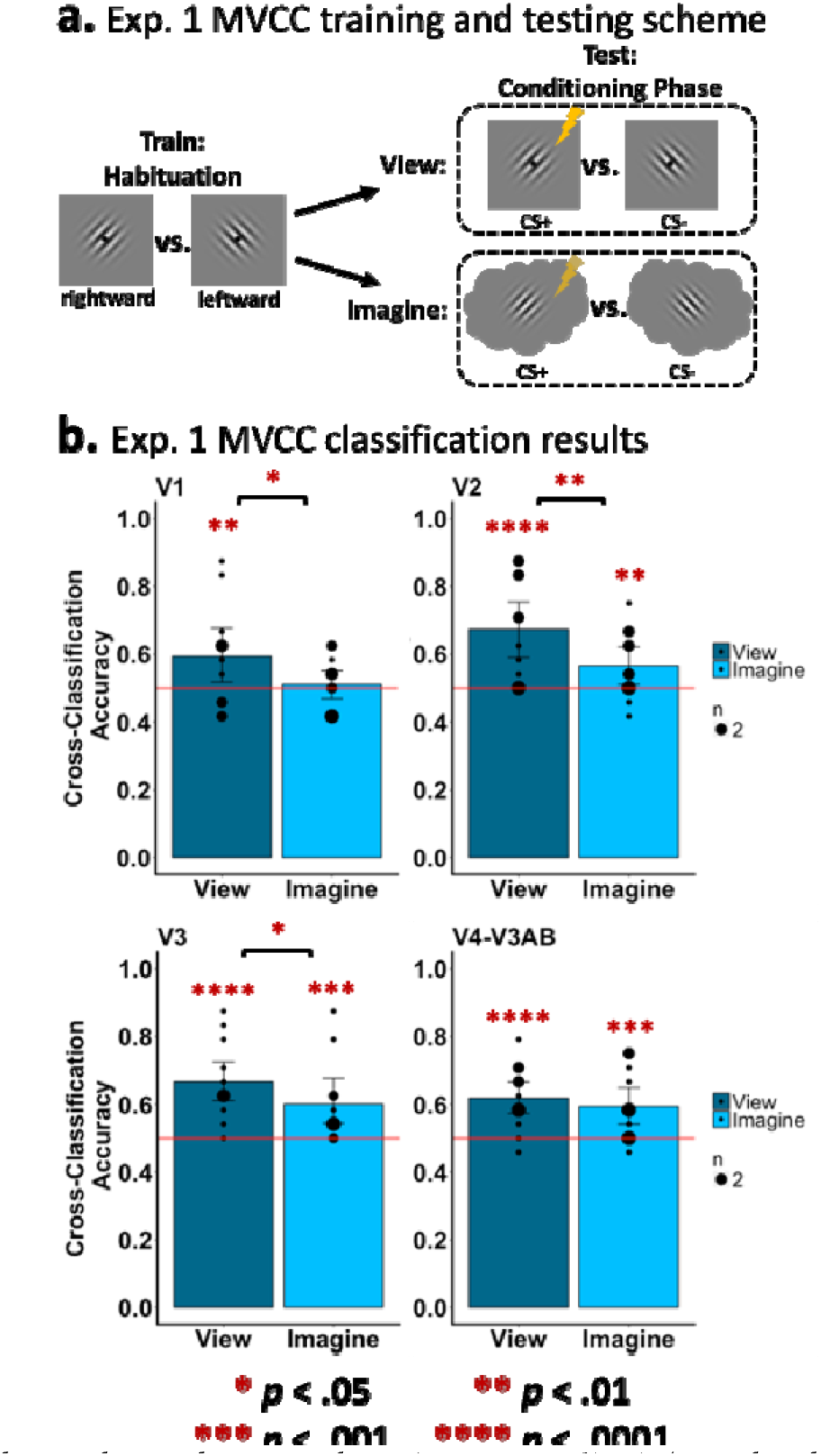
MVCC in early visual cortex during the conditioning phase (experiment 1). a) A graphical representation of the MVCC training and testing scheme. The lightening bold denotes the CS+ but the MVCC analysis excluded trials in which shock was delivered. b) results of the MVCC classification of view trials (vCS+ vs. vCS-; dark-gray bars) and imagine conditions (light-gray bars) in V1 (top-left), V2 (top-right), V3 (bottom-left), and V4-V3AB (bottom-right). The horizontal line represents chance (50%), and the black dots represent how many participants had a given classifier accuracy.

**Figure 3.**
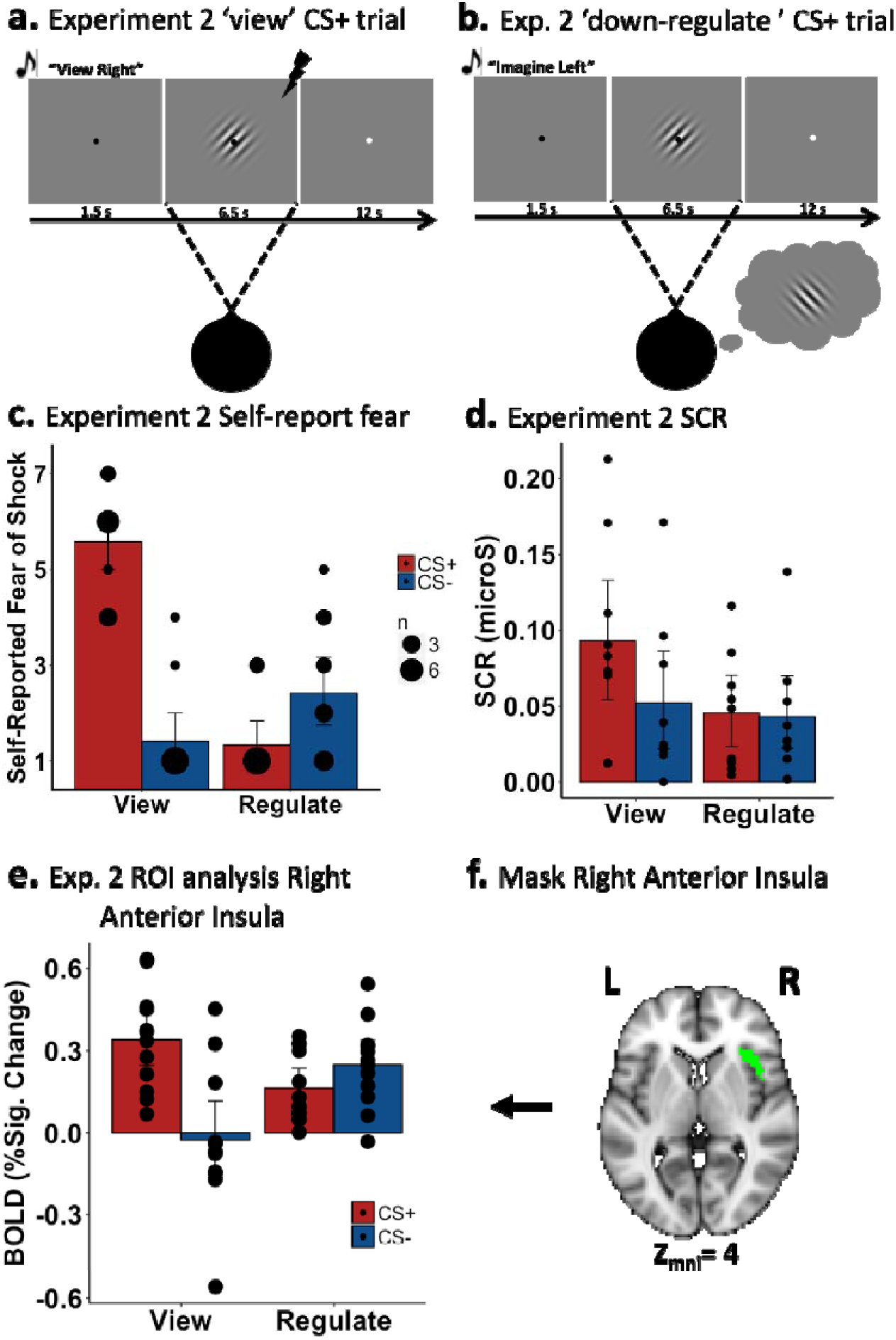
Depiction of the trial sequence for experiment 2 for (a) a ‘view’ CS+ trial, and (b) an ‘down-regulate’ CS+ trial in which participants are cued to imagine the CS- while viewing the CS+. c) Self-reported fear of shock and (d) SCR results. (e) BOLD response for the right anterior insula during experiment 2, emotion regulation phase. (f) The right anterior insula mask was derived from the neural overlap analysis from experiment 1 restricted to the insula. In general, ‘Regulate’ indicates that participants were cued to imagine the opposite stimulus to the one being viewed. Error-bars represent 95% confidence intervals. Black dots the number of participants that shared a given value in the dependent measure.

**Figure 4.**
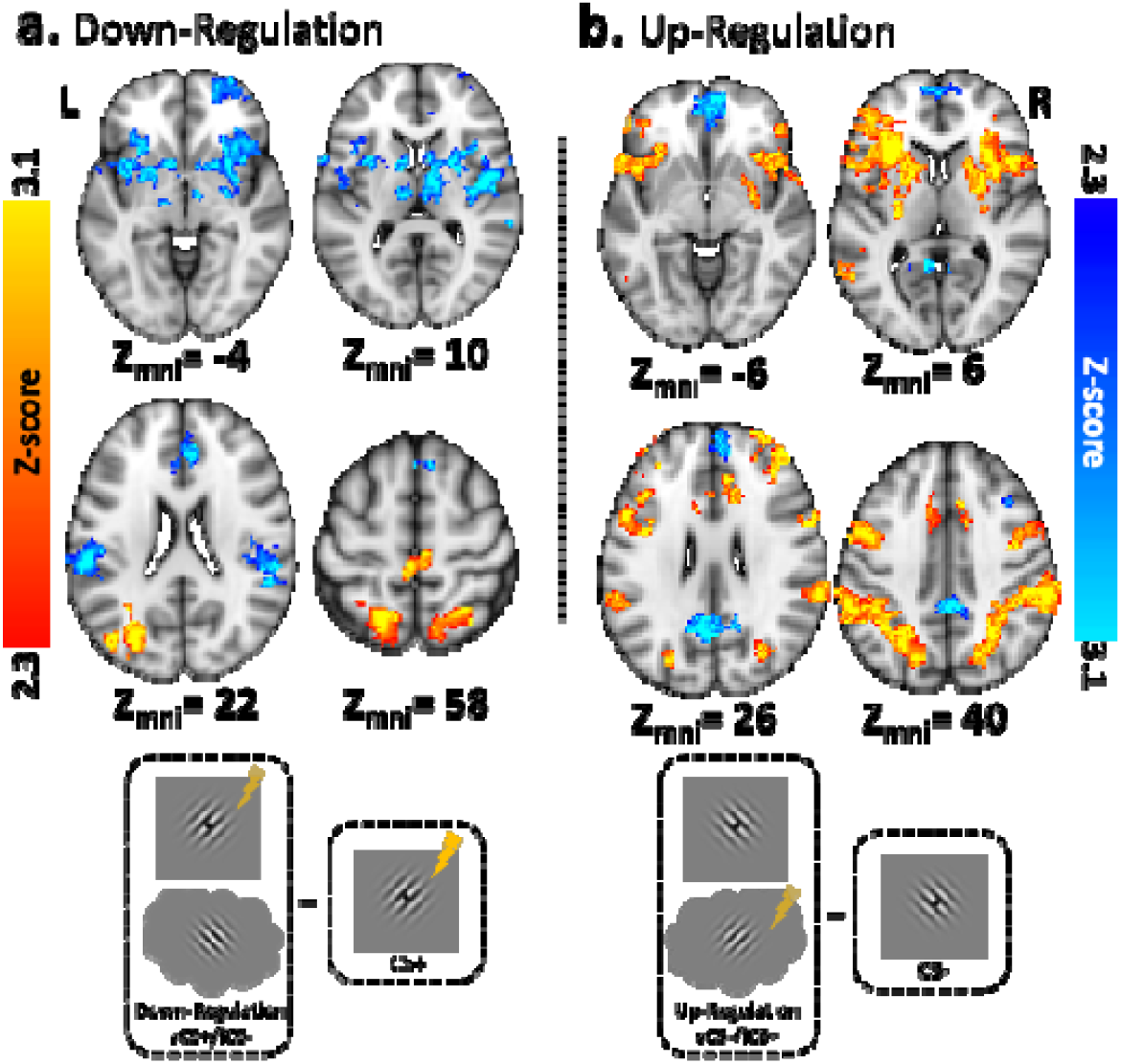
Results from the whole-brain analysis of the emotion regulation phase (experiment 2). a) Results comparing the ‘down regulate’ CS+ (red-yellow) condition to the ‘view’ CS+ (blue-light blue) condition. The ‘down-regulate’ CS+ had participants imagine the CS- while viewing the CS+. b) Results comparing the ‘up-regulate’ CS- (red-yellow) condition to the ‘view’ CS- (blue-light blue) condition. The ‘up-regulate’ CS- had participants imagine the CS+ while viewing the CS-.

**Figure 5.**
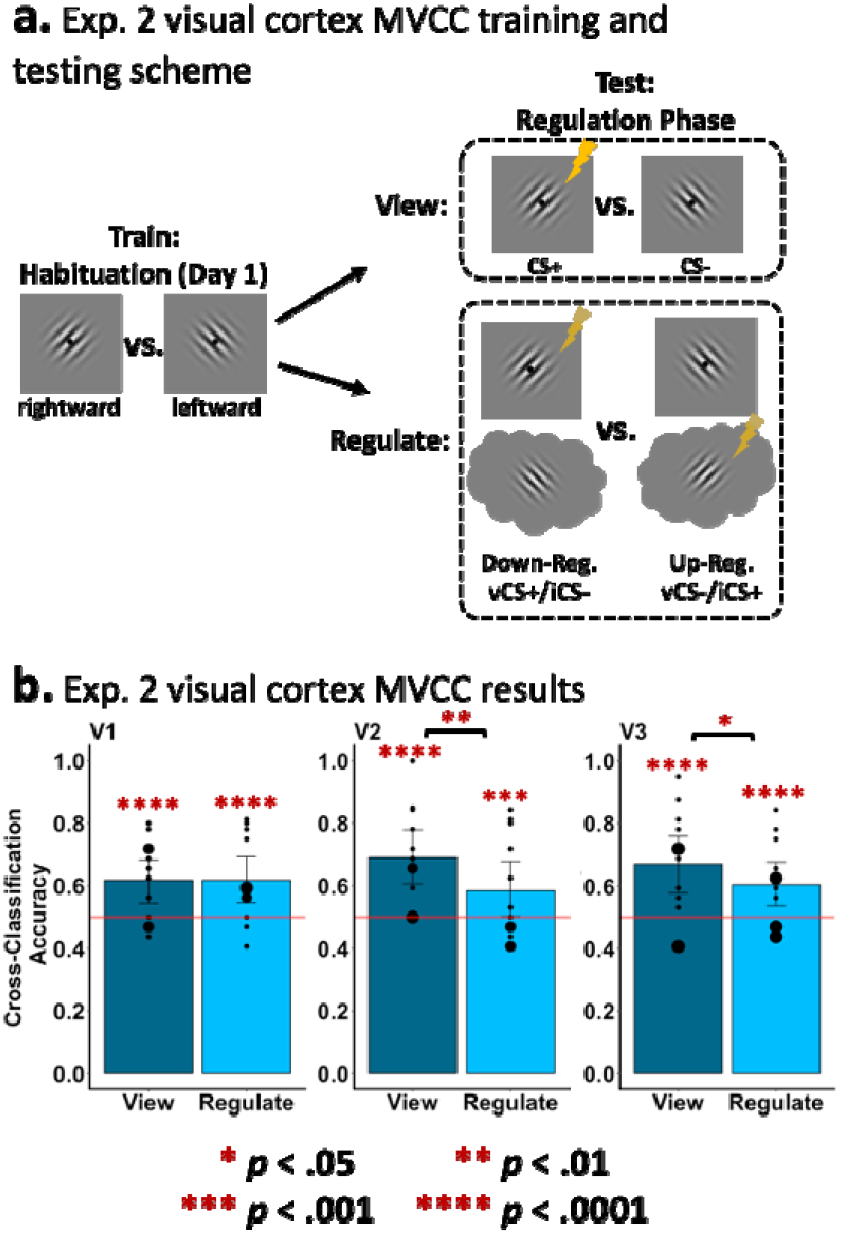
MVCC in early visual cortex during the emotion regulation phase (experiment 2). a) A graphical representation of the MVCC training and testing scheme. The lightening bold denotes the CS+ though the MVCC analysis excluded trials with shock. b) Results of the MVCC classification of view trials (vCS+ vs. vCS-; dark-gray bars) and regulate conditions (‘down-regulate’ (vCS+/iCS-) vs. ‘up-regulate’ (vCS-/iCS+); light-gray bars) in V1 (bottom-left), V2 (bottom-middle), V3 (bottom-right). The horizontal line represents chance (50%), and the black dots represent how many participants had a given classifier accuracy.

**Figure 6.**
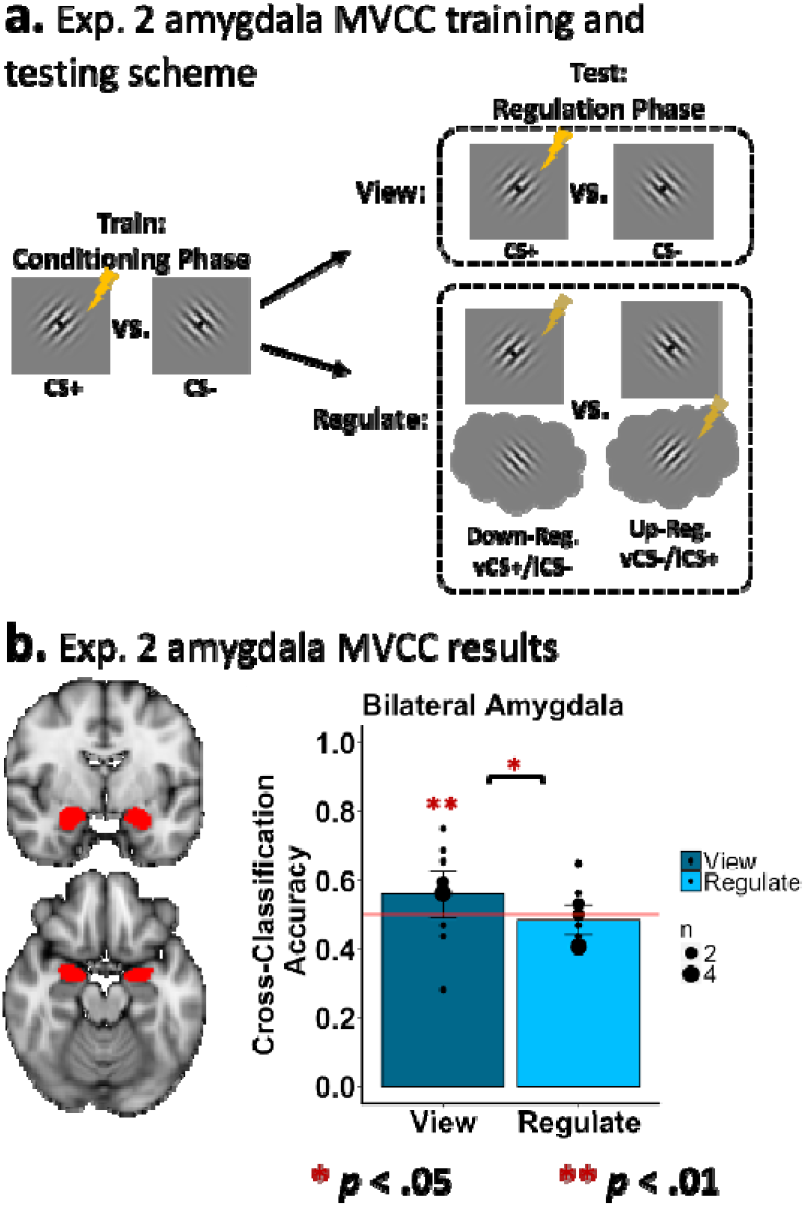
MVCC in the amygdala during the emotion regulation phase (experiment 2). a) A graphical representation of the MVCC training and testing scheme. The lightening bold denotes the CS+. b) Results of the MVCC classification of view trials (vCS+ vs. vCS-; dark-gray bars) and regulate conditions (‘down-regulate’ (vCS+/iCS-) vs. ‘up-regulate’ (vCS-/iCS+); light-gra bars) in bilateral amygdala. The horizontal line represents chance (50%), and the black dots represent how many participants had a given classifier accuracy.

### Conclusion

Taken together, the results of the present study indicate that mental imagery can be cued to successfully generate and regulate a differential fear conditioned association. Furthermore, the findings from MVCC and the early visual cortex are consistent with the depictive theory of mental imagery when imagining a CS+ versus CS- in the absence of external stimuli. Moreover, the MVCC findings that imagining the opposite CS from the one being viewed can regulate emotional reactivity is consistent with both the depictive and biased-competition theories. More broadly, the findings of the present study also support the potential utility of mental imagery-based techniques in the treatment of psychological illness, from imaginal exposure in post-traumatic stress disorder and phobias[81] to imagery rescripting in depression and anxiety[82].

## METHODS

### Participants

Thirteen participants (6 women/7 men, mean age = 24 years, *SD* = 4.87) from the University of Southern California and surrounding community enrolled in this study. All participants provided informed consent and the study was approved by the University of Southern California Institutional Review Board. All procedures were carried out in accordance with the approved guidelines and were in accordance with the ethical principles of the Belmont Report. None had a history of psychological or neurological disorder. Participants completed two experiments across two days of scanning (mean days between sessions = 7, *SD* = 9). One participant dropped out after completing day one (Experiment 1). Thus, Experiment 1 included 13 participants and Experiment 2 included 12. Our sample size was based on other similar studies employing multivariate pattern analysis to study the impact of mental imagery on early visual cortex activity[34,83,84], as well as on studies of fear conditioning using multivariate analysis techniques[49].

### Materials

The conditioned stimuli consisted of two Gabor patches (diameter: 8° visual angle; spatial frequency: 2.1 Hz with randomized spatial phase; contrast ratio: 0.75): one oriented 45° from the horizontal (which was referred to as the ‘rightward’ oriented patch) and one oriented 135° from the horizontal (which was referred to as the ‘leftward’ oriented patch). Whenever a Gabor was presented it flashed on and off at a rate of 2Hz (1,750ms on, 250ms off) with a random spatial phase to avoid adaptation[85]. We chose to use Gabor stimuli as previous research demonstrated that they are sensitive to the effects of emotion[86] and they can be decoded from early visual cortex when being imagined[33]. The unconditioned stimulus (US) consisted of a 500ms (at 50Hz) mild electric shock delivered to the fingertips of the ring and pinky fingers of the left hand[13], administered using E13-22 (Coulbourn Instruments, Allentown, PA), and included MR-compatible leads and electrodes (BIOPAC systems, CA), and a grounded RF filter. The schedule of stimulus presentation was delivered with Psychtoolbox-3[87, 88] in MATLAB (MathWorks, Natick, MA, USA). The auditory instructions were created from www.fromtexttospeech.com (language: US English, Voice: Heather, Speed: medium) and delivered using Sensimetric MRI-compatible Insert Earphones[89].

### Design and Procedure

In the present study we used fear conditioning to study emotional reactivity as it has advantages over other options such as emotional pictures or scenes. For example, while pictures can elicit highly varied and idiosyncratic emotional responses[90, 91], fear conditioning has a higher degree of experimental control and can be confirmed using multiple convergent dependent measures[92, 93]. Fear conditioning also allows for the comparative, cross-species, study of emotional reactivity[15, 94]. In addition, as the focus of the present study is on the multivariate decoding of mental imagery in the early visual cortices it was important that we used conditioned stimuli that were well controlled and did not differ in terms of visual features (e.g, luminance, contrast, colour) and stimulus complexity.

#### Visit 1, Experiment 1 – Fear transfer to imagined stimuli

The first experiment sought to determine if fear conditioning to a visual object transfer to instances of mentally imagining the fear conditioned stimulus. This tests the prediction that mental imagery of a CS+ versus CS- generates differential fear response. At the beginning of Experiment 1, electrodes for electrical stimulations were secured and the shock level was adjusted individually to be “unpleasant but not painful” (*M_intensity_* = 2.08 mA, *SD* = 0.67, range: 1.10-4.00 mA). Once in the MRI scanner, participants performed three consecutive phases: the classifier training phase in which participants only viewed the CSs (i.e., there was no mental imagery); the pre-conditioning phase in which participants both viewed and imagined the CSs; and the conditioning phase in which participants viewed and imagined the CSs. The CS+ was paired with shock on 50% of view trials during the conditioning phase.

The classifier training phase included six runs composed of five visual presentations of each Gabor patch (i.e., the leftward and rightward oriented patches) in a random order with no audio, resulting in 30 trials per patch across the six runs. Each trial began with a Gabor patch presented for 6.5s and ended with a 12s inter-trial interval (ITI). The slow-event trial structure used in this phase and all the others was modelled on the fear conditioning and regulation study by Delgado et al.[9]. This offered a structure similar to critical studies that decoded imagery of visual objects in early visual cortices, which used slow-event designs and did not jitter the ITI[33,63,84]. Additionally, on the recommendation of Coutanche and Thompson-Schill[95], we used multiple shorter runs in this and every other phase of the experiment to improve the generalizability of our MVCC classification models. Participants were instructed to maintain fixation on a center fixation dot (0.5° visual angle), which turned black when a Gabor was present and white during the ITI. The purpose of the classifier training phase was to provide training data to our classification model such that it could accurately decode the Gabor patches (i.e., the rightward versus leftward patch) in early visual cortices during the two critical experimental phases, the fear conditioning phase (Day 1) and the regulation phase on Day 2 (see the MVCC description below). Notably, this involved presenting the visual Gabor patches before any conditioning or imagery took place so that the classifier was not confounded by these extraneous factors.

Following classifier training, participants completed the pre-conditioning phase which comprised three runs that were each composed of four ‘view’ trials and four ‘imagine’ trials of each Gabor patch, for a total of 12 trials per condition across the three runs. The purpose of the pre-conditioning phase was to ensure participants practiced following the instructions including the use of instructed mental imagery. At the beginning of the pre-conditioning phase participants were reminded to maintain fixation on the center dot, and they were instructed “when the fixation dot turns black this signals the start of a trial and you should listen for and follow the instructions; when the fixation dot turns white the trial is over and you can relax and prepare for the next trial”. During pre-conditioning, each trial began with an audio instruction (1.5s), which was one of four possibilities: “view right”, “view left”, “imagine right”, “imagine left”. Next, either one of the two Gabor patches appeared along with the fixation (following “view right” or “view left” instructions) or the fixation appeared alone for 6.5s (following “imagine right” or “imagine left” instructions). The trial ended with a 12s ITI. The trial presentation order was random. There were no trials in which a participant was cued to “imagine” followed by the visual presentation of a Gabor patch, nor were there trials in which a cue to “view” was followed by the fixation alone as on typical imagery trials.

Most importantly, the final phase of Visit 1 was the fear conditioning phase, which comprised six runs composed of four ‘view’ trials and two ‘imagine’ trials of each Gabor patch, for a total of 12 trials per run (see Figures 1a & 1b for a depiction of a view CS+ trial and an imagine CS+ trial, respectively). The general trial structure was identical to the structure of the pre-conditioning phase with the notable exception that one of the two Gabor patches (the order was counterbalanced across participants) was designated as the CS+ and co-terminated with shock on 50% of ‘view’ CS+ trials. The second Gabor patch, designated as CS-, was never associated with shock. No shock was ever delivered on ‘imagine’ trials. In addition, the first and the last trial of each run was a ‘view’ CS-. The second trial of each run was always a ‘view’ CS+ with shock. All the remaining trials (from trial 3 to 11) were randomized. As with previous research[9, 13], we excluded the two reinforced CS+ trials of each run from the analyses. We also excluded the first and last CS- trials of each run. This ensured that the number trials per condition included in the analyses were equal and controlled for potentially spurious novelty effects generated on the first trial of each run[13]. Overall, this produced 12 total trials per condition across six runs for inclusion in our analyses.

Due to time constraints at the scanner one participant completed 5 classifier training runs (25 trials per condition), 2 pre-conditioning runs (8 trials per condition) and 3 conditioning runs (6 trials per condition), while a second participant completed only 2 pre-conditioning runs (all other runs were completed).

#### Visit 2, Experiment 2 – Regulation of fear conditioning via mental imagery

In the second experiment, we sought to determine if mental imagery can exert an emotion regulation effect such that an internally generated representation competes with an externally generated representation, one of which involves emotional content. Experiment 2 was completed on day two and involved a similar set-up to Experiment 1. First, the electrodes for electrical stimulations were secured and the shock level was adjusted individually to be “unpleasant but not painful” (*M_intensity_* = 2.00 mA, *SD* = 0.32, range: 1.40-2.30 mA).

Once in the scanner, participants completed the emotion regulation phase, eight runs that each contained four ‘view’ trials and two ‘regulate’ trials of each CS for a total of 12 trials per run. The general task structure was identical to the conditioning phase of Visit 1, except the ‘imagine’ trials of Visit 1 were replaced with ‘regulate’ trials (see Figure 3a & 3b for a depiction of a ‘view’ CS+ and a ‘down-regulate’ CS+ trial, respectively). The CS+ was the same Gabor across Experiment 1 and Experiment 2 for each participant, as was the CS-. The ‘view’ trials began with the audio instruction to either ‘view right’ or ‘view left’, which was followed by the matching Gabor. The ‘regulate’ trials began with an audio instruction to either “imagine left” or “imagine right”. Next, the physical presentation of a Gabor patch began during which participants had to imagine the instructed Gabor patch. While not told explicitly to participants, during ‘regulate’ trials participants were always instructed to imagine the opposite Gabor from the one presented visually. This produced two ‘regulate’ conditions: The ‘down-regulation’ condition during which participants viewed the CS+ while imagining the CS- (vCS+/iCS-); and, the ‘up-regulation’ condition during which participants viewed the CS- while imagining the CS+ (vCS-/iCS+). No shock was delivered on ‘regulate’ trials. Fifty percent of ‘view’ CS+ trials were reinforced with a co-terminating shock. We used the same approach as in the conditioning phase with regards to trial presentation order, with a ‘view’ CS- for the first and the last trial of each run, and a ‘view’ CS+ with shock for the second trial. All the remaining trials (from trial 3 to trial 11) were randomized in each run. Moreover, as with the conditioning phase, all reinforced CS+ trials were excluded from the analyses along with the first and last CS- trials. Overall, this produced 16 total trials per condition across six runs for inclusion in our analyses. Due to time constraints at the scanner one participant completed only 5 of the 8 regulation phase runs (i.e., 10 trials per condition). During all experimental stages across both days, the participants were attached to the SCR and shock electrodes, and the shock stimulator was set to the ‘On’ position.

### Self-report measures

After each experiment, participants provided self-reported evaluations of: 1) the vividness of their mental images on ‘imagine’ trials, on a 7-point scale ranging from “1 = Non-Existent” to “7 = Very Strong” (e.g., “How vivid was your mental imagery on IMAGINE LEFT trials?”); 2) their effort to form the mental images, on an 7-point scale ranging from “1 = Not At All” to “7 = Very Hard” (e.g., “How hard did you try to form the mental images on IMAGINE RIGHT trials?”); and, 3) their fear of getting shocked on each type of trial, on a 7-point scale ranging from “1 = Not At All” to “7 = Very Much So” (e.g., “How much did you fear the shock on VIEW RIGHT trials?”). For each question participants were asked specifically about one of the two Gabor patches. Moreover, so as to not bias participants no reference was made to the terms “CS+” or “CS-”.

### SCR Methods and Analysis

The SCR collection and analysis were carried out consistent with previous research from our group[96–98]. We measured skin conductance responses (SCRs) during MRI acquisition. The physiological data was recorded at a sampling rate of 1 kHz using BIOPAC’s MP-150 system (BIOPAC System, Goleta, CA, USA). We employed grounded RF filtered MR-compatible leads and MRI-compatible Ag/AgCl electrodes placed on the fingertips of the index and middle fingers of participants’ left hand. Trials including shock delivery were excluded from all analyses, though for the first level fMRI analyses these trials were included as nuisance regressors. Additionally, the first and last trial of each run was a CS- trial which was excluded from the analysis, though they were also included as nuisance regressors for fMRI analyses. This was done to eliminate the potential confounding orienting response to the first trial and to ensure that each condition in the primary analyses included an equal number of trials[13]. Offline, the SCR data were detrended and smoothed with a median filter over 50 samples to filter out MRI-induced noise and down-sampled to 100 Hz[13]. On a trial-by-trial basis, the SCR epochs were extracted from a time window between 0 and 8 s after CS onset and baseline-corrected by subtracting the mean signal from the one second before CS onset[98]. Next, the SCR peak was calculated by taking the maximum SCR response from 1-8 s and subtracting the minimum SCR value from 0-.99 s on a trial-by-trial basis, and any response less than 0.02 μS was zeroed. Finally, the resulting SCR peak data averaged over each condition per participant. One participant had no detectable SCR response to the US and was excluded from all SCR analyses, and the SCR equipment malfunctioned during fMRI for two participants during the entire regulation phase. Additional SCR malfunctions occurred for 2 participants on a subset of the data: One participant had no data for one run of the conditioning phase and 2 runs of the regulation phase; the second participant had no data for one run of the regulation phase. Finally, one participant did not complete all runs, as noted above, and completed only 3 conditioning runs (6 trials per condition) and 5 regulation runs (i.e., 10 trials per condition). All data that we did collect for these participants were retained and included in the SCR analyses.

In Experiment 1, following our repeated measure analysis of variance (ANOVA) on the SCR data we conducted one-tail tests on our *a priori* predictions regarding a larger SCR response to the CS+ versus CS- for each the view and imagery conditions. One-tailed tests were used because we had a specific directional hypothesis regarding SCR and fear conditioning, which is also theoretically meaningful. In the case of Experiment 1, it is of no theoretical importance to consider that the CS- might elicited a greater SCR than the CS+. This consistent previous approaches by us[97] and others[11,28,99] in studies involving human fear conditioning. On the other hand, in Experiment 2 we used two-tailed tests as it was theoretically meaningful particularly for comparing the two regulate conditions, the ‘regulate’ CS+ condition (vCS+/iCS-) versus the ‘regulate’ CS- condition (vCS-/iCS+).

### MRI Data Acquisition and Analysis

#### MRI acquisition

Imaging was performed using a 3T Siemens MAGNETON Trio System with a 32-channel matrix head coil at the Dana and David Dornsife Neuroscience Institute at the University of Southern California. A T1-weighted high-resolution image was acquired on both Visit 1 and Visit 2 of scanning using a three-dimensional magnetization-prepared rapid acquisition gradient (MPRAGE) sequence (TR = 2530 msec, TE = 3.13 msec, flip angle = 10°, 224 × 256 matrix, phase encoding direction right to left). 176 coronal slices covering the entire brain were acquired in sequential order with a voxel resolution of 1mm isotropic.

All functional images (with the exception of the retinotopic mapping) were acquired using a gradient-echo, echo-planar, T2*-weighted pulse sequence (TR = 2000 msec, TE = 25 msec, flip angle = 90°, 64 × 64 matrix, phase encoding direction posterior to anterior). Thirty-eight slices covering the entire brain were acquired with an in-plane voxel resolution of 3.0 × 3.0 and a slice thickness of 3 mm with no gap. Slices were acquired in interleaved ascending order, and each functional run began with the collection of 4 dummy volumes to account for T1 equilibrium effects, which were discarded as part of the later preprocessing steps of data analyses. The total number of volumes per run, including dummy volumes, varied according to the experimental phase: Classifier Training (Visit 1) = 98 volumes, Pre-Conditioning (Visit 1) = 85 volumes, Conditioning (Visit 1) = 125 volumes, and Regulation (Visit 2) = 125 volumes.

At the end of Visit 2 we performed two functional retinotopic mapping scans, one for polar angle and one for eccentricity. They were acquired using a gradient-echo, echo-planar, T2*-weighted pulse sequence (TR = 1,200 msec, TE = 30 msec, flip angle = 65°, 78 × 78 matrix, phase encoding direction posterior to anterior). Twenty slices covering the occipital lobe and positioned perpendicular to the calcarine sulcus were acquired with an in-plane voxel resolution of 2.5 × 2.5 and a slice thickness of 2.5 mm with no gap. For each, 254 total volumes were collected including 4 dummy volumes.

We also collected a T2-weighted anatomical scan on Visit 1 (TR = 10,000 ms, TE = 88 ms, flip angle = 120°, 256 × 256 matrix) with 40 transverse slices with a voxel resolution of 0.82 × 0.82 × 3.5 mm. This scan was reviewed by a radiologist to rule out incidental findings and ensure all participants were neurologically normal.

#### fMRI Preprocessing and Whole-brain Univariate Analysis

FMRI data processing was carried out using FEAT (FMRI Expert Analysis Tool) Version 6.00, part of FSL (FMRIB’s Software Library, www.fmrib.ox.ac.uk/fsl). Registration of the functional images to both the high resolution (T1-weighted) structural image and the standard space image was carried out using FLIRT[100, 101]. The following pre-statistics processing was applied: motion correction using MCFLIRT[101]; slice-timing correction using Fourier-space time-series phase-shifting; non-brain removal using BET[102]; spatial smoothing using a Gaussian kernel of FWHM 5mm; grand-mean intensity normalization of the entire 4D dataset by a single multiplicative factor; high-pass temporal filtering (Gaussian-weighted least-squares straight line fitting, with sigma=50.0s). The time-series statistical analysis was carried out using FILM with local autocorrelation correction[103].

The data were analyzed within the General Linear Model using a multi-level mixed-effects design. At the single-participant level each run was modelled separately. We used a double-gamma hemodynamic response function (HRF) with which each of the conditions of interest (i.e., onset to offset of either viewing or imaging a given Gabor) was convolved. We included a model for the temporal derivative of each condition of interest. We also included several nuisance regressors including six motion correction parameters, and motion censoring regressors for any volume with >0.9mm framewise displacement[104] using the fsl_motion_outliers function. As noted previously we also modelled CS+ reinforced (i.e., shock) trials and the first and last CS- trials as two separate regressors that were not used in higher-level analyses (they were modelled similar to the conditions of interest). A second-level analysis was performed in order to combine contrast estimates from the first level separately for each experimental phase (e.g, the conditioning phase) for each participant. This was completed using a fixed effects model, by forcing the random effects variance to zero in FLAME (FMRIB’s Local Analysis of Mixed Effects)[105–107]. Group-level analyses were carried out using FLAME (FMRIB’s Local Analysis of Mixed Effects) stage 1 and stage 2 with automatic outlier detection[105–107]. The resulting Z (Gaussianised T/F) statistic images were corrected for multiple comparisons using FSL’s cluster thresholding algorithm that applies Gaussian Random Field Theory to estimate the probability of observing clusters of a given size. We applied a threshold of Z>2.3 and a (corrected) cluster size probability of *p*=0.05[108]. All whole-brain unthresholded group-level maps can be viewed at neurovault.org using the link: https://identifiers.org/neurovault.collection:5138

#### Neural Overlap Analysis

All references to the neural overlap analysis involved the quantification of the extent of neural overlap between two independent whole-brain group-level maps both of which will have been previously thresholded (*z* > 2.3) and corrected for multiple comparisons[109, 110]. To quantify the extent of neural overlap when experiencing a differential fear conditioned response to viewed stimuli compared to imagined, we performed a neural overlap analysis on whole-brain group-level maps from the ‘view’ CS+ > ‘view’ CS- analysis and the ‘imagine’ CS+ > ‘imagine’ CS- analysis (Conditioning phase, Experiment 1).

#### Context-Dependent Functional Connectivity (Psychophysiological Interaction, PPI)

To assess differences in task-specific correlations between a seed region (right aIn) and other areas we performed a psychophysiological interaction (PPI) analysis[111] on the data from both the conditioning phase (Visit 1) and the regulation phase (Visit 2). In both cases the seed region was produced from the right anterior insula cluster that resulted from the neural overlap analysis of the conditioning phase. This cluster was further restricted by right insula mask of the Harvard-Oxford Cortical atlas. This analysis was performed to test the prediction that modulation of aIn occurred via distinct neural pathways in different conditions. Specifically, we predicted that whereas ‘view’ conditions would involve greater connectivity with bottom-up areas such early visual areas and the thalamus, conditions involving imagery (‘imagine’ CS+ on Visit 1, and ‘down-regulate’ CS+ on Visit 2) would involve greater connectivity with regions associated with cognitive control and memory such as frontoparietal and medial temporal lobe (MTL) regions, respectively.

At the single-participant (first-level) we ran a general linear model (GLM) that included: our psychological regressor contrasting our two conditions of interest [i.e., Visit 1: (‘view’ CS+) - (‘imagine’ CS+)); Visit 2: (‘view’ CS+) - (‘down-regulate’ CS+)]; our physiological regressor, which was the time-series of the right aIn extracted from the preprocessed and filtered data supplied to our initial first-level analyses (with no convolution, no temporal derivative and no temporal filtering); and the critical PPI regressor, which is modelled as the interaction between the first two regressor such that the physiological regressor zero centered on the mean (i.e., the ‘mean’ option) and the psychological regressor is zero centered on the halfway point between the highest and lowest point of the regressor (i.e., the ‘center’ option). No temporal derivative or temporal filtering was applied to the PPI regressor. Our PPI modelled also included nuisance regressors which included one regressor reflecting the shared variance of the two PPI conditions of interest [Visit 1: (‘view’ CS+) + (‘imagine’ CS+)); Visit 2: (‘view’ CS+) + (‘down-regulate’ CS+)], all the other conditions, six motion correction parameters, and motion censoring regressors (the same as described in the main level 1 analysis).

#### Functional Retinotopic Localizer

At the end of day two, all participants completed two functional retinotopic localizer runs: one polar angle, and one eccentricity run[112, 113]. Both functional localizers involved a flickering checkerboard (5Hz), the polar angel localizer used a rotating wedge and the eccentricity localizer used an expanded annulus. Each localizer run involved ten cycles, with 25 fMRI volumes per cycle (volumes were 1.2 seconds). The maximum visual angle reach by both the wedge and the annulus was ∼8.4 degrees. Participants were instructed to keep their overt attention on the center fixation and to track covertly the flickering checkerboard for randomly presented circular targets. The target had a 5% chance of occurring with each flicker and was presented for 200ms. Participants reported the number of targets they detected at the end of each run orally, however, this information was not record as it was simply to ensure participants were attentive. The primary aim of the retinotopic mapping was to produce individual ROIs of both the bilateral dorsal and ventral aspects of each V1, V2, V3, and V4-V3AB combined[114, 115].

Functional retinotopy data was preprocessed using Freesurfer’s standard procedures for polar retinotopic mapping, including spatial smoothing with 5mm kernel, resulting in a flattened cortical surface[116, 117] that was cut along the calcarine sulcus (http://freesurfer.net/fswiki/FreeSurferOccipitalFlattenedPatch). The functional data were overlapped on the flattened map and ROIs were manually traced individually for left and right hemispheres and for the ventral and dorsal aspects of each visual area. After tracing, the surface-based ROIs were converted to volumetric space using Freesurfer then registered to the native space of each participants’ functional experimental data using FSL’s FNIRT (FMRIB’s Nonlinear Image Registration tool) with trilinear interpolation[118]. The resulting ROIs were thresholded (50%) and visually inspected to ensure no overlap between ROIs.

#### Multivariate pattern analysis and cross classification in visual cortex

We used MVCC[48] to evaluate critical aspects of both the conditioning phase and the regulation phase. MVCC involves training a machine learning classifier algorithm in one context or condition and evaluating the performance of the classifier in a different context or condition. If there is significant cross-classification performance this provides evidence of informational similarity between the contexts or conditions. Relevant to the current study, previous research has demonstrated that a classifier trained to discriminate between visual objects such as line patches or letters, can be used to quantify participants degree of visual imagery[33,34,63] or the focus of their visual attention[85,119,120].

For the conditioning phase, we used MVCC to identify whether when participants were asked to imagine a given Gabor they did so (Figure 2). The outcome of this analysis could provide evidence that generating a decodable signal in early visual cortex is associated with generating a fear conditioned response. For the regulation phase, MVCC allowed us to test the prediction that imagining one stimulus while viewing a second stimulus disrupts the neural representation of each via biased competition. Overall, our prediction is that such competition would result in reduced classifier accuracy during the regulation conditions (during which participants viewed one Gabor while imagining the opposite one) compared to the view conditions.

In the present study, we used the data from our day one classifier training phase to train classifiers on each visual cortex ROI for decoding the ‘leftward’ versus ‘rightward’ Gabor patches, during which fear conditioning had not occurred thus ensuring that the classifier was not confounded by any effects of associative learning (See Figure 2a & 5a for a visual depiction of our MVCC approach in both the conditioning phase and the regulation phase, respectively). Our ROIs were derived from our functional retinotopic localization of V1, V2, V3, and V4-V3AB. All participant specific data and masks used in this analysis were registered to the middle functional volume of the first classifier training run.

Preprocessing and first-level modeling of the fMRI data used for MVCC was identical to the univariate analysis with the notable exception that each trial was modelled independently in the first-level GLM. This included smoothing with a 5mm FWHM kernel, which has been shown previously to improve classifier performance[121]. In addition, the data were analyzed in native space, and were not transformed to standard space. These parameters were chosen following an iterative evaluation using leave-one-run-out cross-validation using only the training set data (i.e, the classifier training phase data). None of the critical test data, which were the data from the conditioning and regulation phases, were used in parameter optimization.

The MVCC analyses were carried out using PyMVPA[122] on each individual participant, then the within-participant results were aggregated to the group-level for analysis (see below). We used support vector machine (SVM) classification using pyMVPA[122]. As is the default in PyMVPA, the SVM hyperparameter ‘C’ was determined via automatic scaling according to the norm of the data. We used a feature selection step prior to classification for each ROI in which we selected the 120 voxels with the largest positive univariate signal difference[33, 63]. For each participant/ROI combination we trained the SVM on classifier training phase data to classify the presence of the CS+ versus the CS- (or more specifically, the Gabor patch that represented each for a given participant) trial-by-trial. Next, this model was applied to our testing data from the conditioning and regulation phases trial-by-trial. This resulted in two classification accuracy measurements during each phase. In the conditioning phase, we had the accuracy for both classifying ‘view’ CS+ versus ‘view’ CS- and for classifying ‘imagine’ CS+ versus ‘imagine’ CS-. In the regulation phase, we had the accuracy for both classifying ‘view’ CS+ versus ‘view’ CS- and for classifying ‘regulate’ CS+ versus ‘regulate’ CS-. In all instances, chance performance for the classifier was 50%.

For each ROI we estimated a group-level null distribution by combining permutation testing and bootstrapping[123, 124]. At the single participant level, we ran 10,000 iterations in which we trained the SVM on data with randomly permuted target labels within the training set (i.e., the classifier training data from Visit 1) and tested on the held-out testing data. This produced a null distribution for each participant. Notably, for a given ROI the null distribution used for evaluating classification accuracy on view trials was made comparable to the null used to evaluate imagine or regulation trials by using a seeded random number generator for permuting the target labels of the test set. This is relevant to the between condition analysis below.

Next, to generate a group-level null distribution we used a bootstrapping procedure with 10,000 iterations in which we iteratively generated group mean accuracy estimates by randomly sampling from each participants null distribution with replacement at each iteration (i.e., during each iteration we randomly selected one element from each participant’s previously computed null distribution with replacement). Finally, to assess the between condition (e.g., ‘view’ vs. ‘imagine’ trials) classification accuracy in a given ROI we compared the accuracy difference between the empirical data to a null distribution of the accuracy differences from the permuted data (10,000 iterations). We also verified that chance performance for all simulation tests was ∼50%, which further confirms that there was no systematic bias in our classification.

#### MVCC analysis of the bilateral amygdala

Although the meta-analytic evidence indicates that there is no reliable univariate response in the amygdala to differential fear conditioning[6], previous research has found that CS+ versus CS- conditions can be decoded in the amygdala using multivariate pattern analysis[49]. A bilateral amygdala mask from the Harvard-Oxford atlas (thresholded at 50%) was registered to each subject’s native space in order to conducted trial-wise within-subject cross classification. Consistent with Bach et al.[49], 300 voxels with the largest positive univariate signal difference were selected as the features of the model at the subject level. All other MVCC parameters and analysis approaches were as described in the previous section.

To decode CS+ versus CS- during the regulation phase we trained the classifier on data from the day one conditioning phase. In doing so, we first tested the classifier on ‘view’ CS+ versus ‘view’ CS- trials during the regulation phase as a positive control. After observing successful cross-classification in this positive control we tested the classifier on ‘down-regulate’ CS+ versus ‘up-regulate’ CS- trials. This allowed us to test the prediction that mental imagery regulation would significantly reduce classification accuracy of patterns associated with differential fear conditioning. We also trained a model on the ‘view’ CS+ versus ‘view’ CS- trials from the regulation phase (Visit 2) to decode CS+ versus CS- during the conditioning phase (Visit 1). However, the positive control of testing the classifier on ‘view’ trials during the conditioning phase was not successful (*p* = .23), thus we were not justified in testing this classifier on imagery trials during acquisition.

## DATA AVAILABILITY

The unthresholded brain imaging results from the univariate fMRI analyses can be found hosted on neurovault.org, using https://identifiers.org/neurovault.collection:5138.

The behavioural data from the current study are available at https://osf.io/r87z9/?view_only=5467316afe3940deb10d81792f3bfb52. The MRI dataset from which the presented findings were generated will be made available at OpenNeuro.org, using 10.18112/openneuro.ds003425.v1.0.0.

## CODE AVAILABILITY

The code associated with the experimental tasks are publicly available at https://osf.io/r87z9/?view_only=5467316afe3940deb10d81792f3bfb52.

## Acknowledgements

This research was supported by University of Southern California Postdoctoral Scholars Research Grant and a Louisiana Board of Regents – Research Competitiveness Subprogram grant to S.G.G.

The authors would like to thank Dr. Sam Schwarzkopf for providing the functional retinotopy localizer scripts.

