## Supplemental for "Mental imagery can generate and regulate acquired differential fear conditioned reactivity"

**Fear in the mind’s eye:**

Steven G. Greening et al.

**Experiment 1 (Visit 1) – Fear generalization to imagined stimuli**

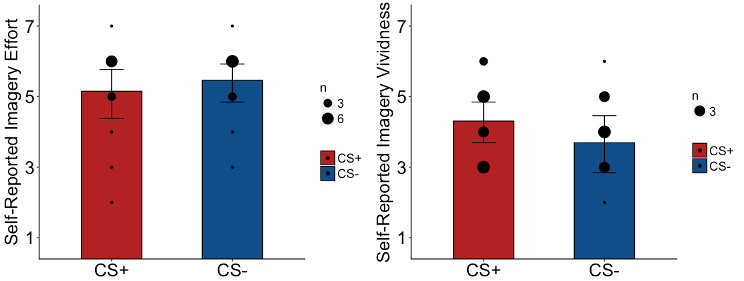

FIGURE S1. Experiment 1 Post-task questionnaire. LEFT – Participants rated their imagery effort, from 1 “Not At All” to 7 “Very Hard”. For example, participants were asked “How hard did you try to form the mental images on IMAGINE RIGHT trials?”. There was no significant differences in imagery effort between imagining the CS+ versus CS- [*t*(12)=1.00, *p* > .3]. RIGHT – Participants rated their imagery vividness, from 1 “None Existent” to 7 “Very Strong”. For example, participants were asked “1)	How vivid was your mental imagery on IMAGINE RIGHT trials? There was no significant difference in self-reported imagery vividness for CS+ versus CS- trials [*t*(12)=1.38, *p* > .19]. Error-bars represent 95% confidence intervals. Black dots represent individual data points, the size of which represents the number of participants that endorsed a given response.

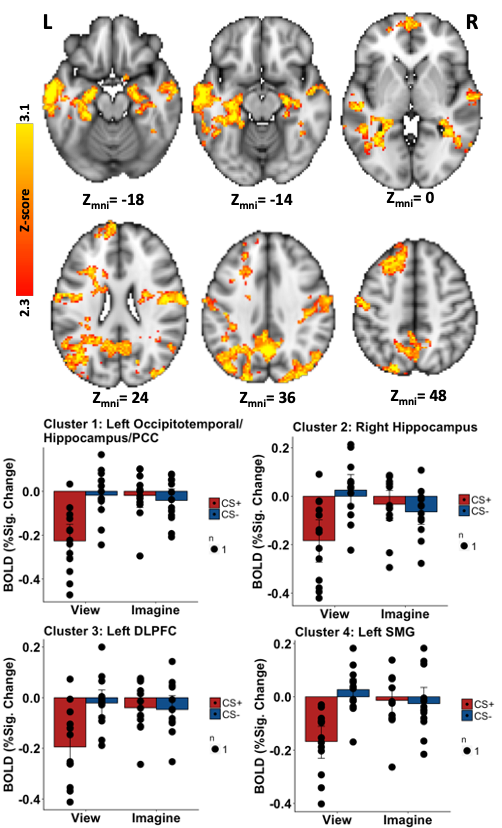

FIGURE S2. Experiment 1 BOLD response for whole-brain interaction analysis (iCS+ > iCS-) > (vCS+ > vCS-). TOP – Activation maps from the whole brain analysis. All clusters were thresholded at z=2.3 and whole-brain corrected for multiple comparisons. Active clusters are displayed on the MNI-2mm standard brain. BOTTOM – Bar graphs of the mean BOLD activation (y-axis) from the significant clusters identified in the whole-brain interaction (see TOP). The graphs allow for qualitative visualization of the interaction effect.

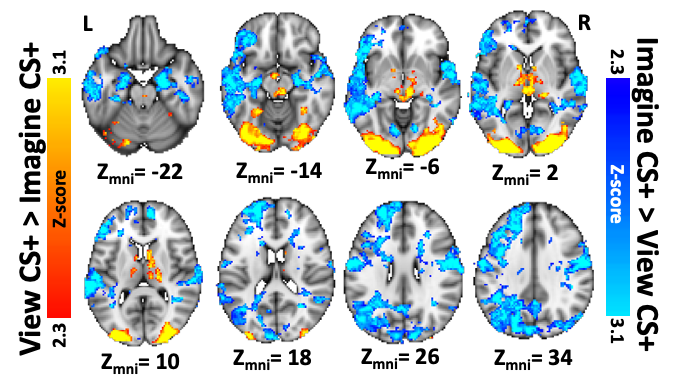

FIGURE S3. Experiment 1 BOLD response for the whole-brain analysis comparing activation when viewing versus imagining the CS+ (Red-Yellow: vCS+ > iCS+; Blue-Light Blue: iCS+ > v CS+). All clusters were threshold at z=2.3 and whole-brain corrected for multiple comparisons. Active clusters are displayed on the MNI-2mm standard brain.

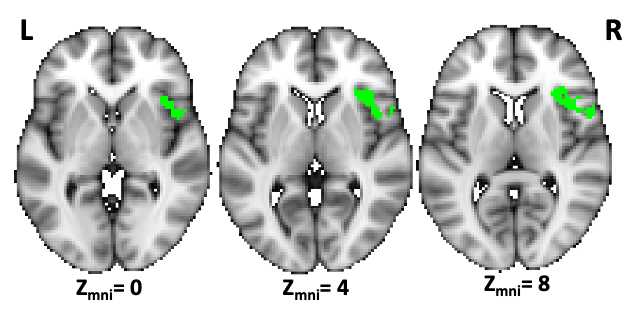

FIGURE S4. Neural Overlap of the thresholded and corrected whole-brain maps from (vCS+ > vCS-) and (iCS+ > iCS-) reveals that both involved activation of the right aIn. The cluster is displayed on the MNI-2mm standard brain.

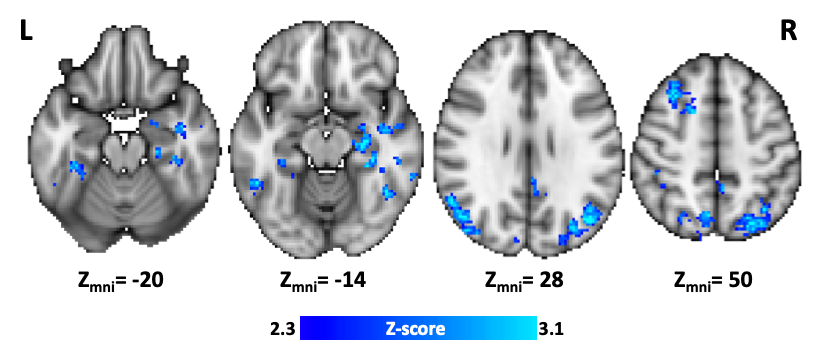

FIGURE S5. Experiment 1 Psycho-physiological interaction (PPI) whole-brain analysis. The seed for the PPI was created by taking the right aIn cluster produced by the spatial conjunction (see Figure S4) and excluding any voxels that were not in the insula (defined as the Harvard-Oxford right insula map). When imagining the CS+ compared to viewing the CS+, greater functional connectivity between the aIN and bilateral aspects of the medial temporal lobe, including parts of the amygdala and hippocampus, as well as bilateral aspects of the inferior parietal lobes and left middle frontal gyrus was observed.

**Experiment 2 (Visit 2) – Regulation of fear conditioning via mental imagery**

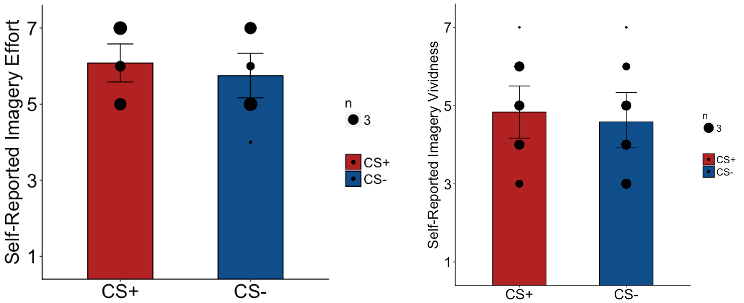

FIGURE S6. Experiment 2 Post-task questionnaire. LEFT – Participants rated their imagery effort, from 1 “Not At All” to 7 “Very Hard”. For example, participants were asked “How hard did you try to form the mental images on IMAGINE RIGHT trials?”. There was no significant differences in imagery effort between imagining the CS+ versus CS- [*t*(11)=1.48, *p* > .16]. RIGHT – Participants rated their imagery vividness, from 1 “None Existent” to 7 “Very Strong”. For example, participants were asked “1) How vivid was your mental imagery on IMAGINE RIGHT trials? There was no significant difference in self-reported imagery vividness for CS+ versus CS- trials [*t*(11)=1.15, *p* > .28]. Error-bars represent 95% confidence intervals. Black dots represent individual data points, the size of which represents the number of participants that endorsed a given response.

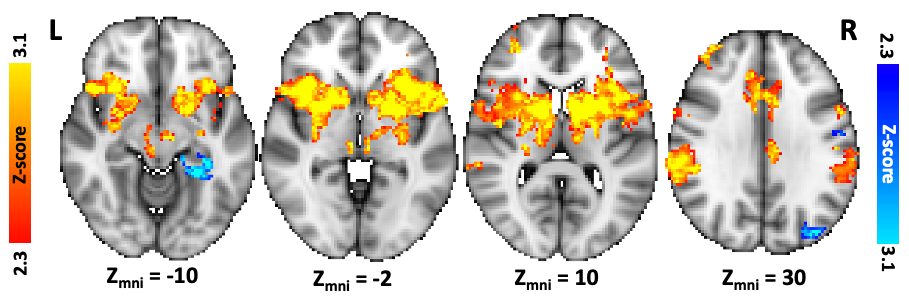

FIGURE S7. Experiment 2 BOLD response for a positive control whole-brain analysis of ‘view’ CS+ versus ‘view’ CS-. Activation maps from the whole brain analysis reveal robust activation of parts of the fear network for CS+ > CS- trials (red-yellow scale) including in bilateral aIn, dACC, thalamus and midbrain areas. There were also several areas more active for CS- > CS+ (blue-light blue) including the right hippocampus. All clusters were threshold at z=2.3 and whole-brain corrected for multiple comparisons. Active clusters are displayed on the MNI-2mm standard brain.

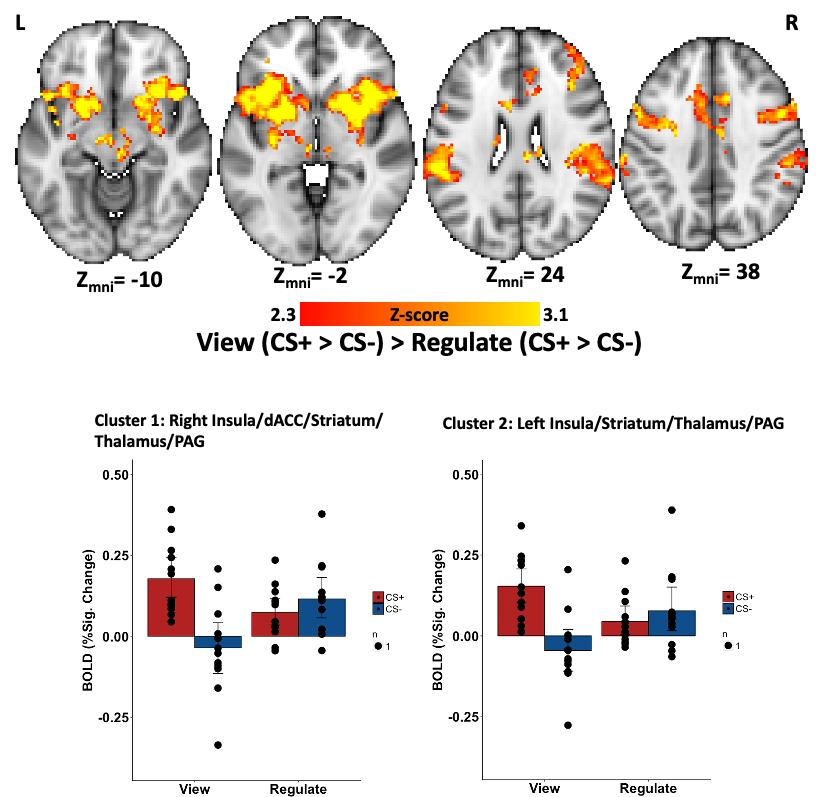

FIGURE S8. Experiment 2 interaction of Stimulus Type (CS+, CS-) by Instruction (View, Regulate). TOP – Activation maps from the whole brain analysis. All clusters were thresholded at z=2.3 and whole-brain corrected for multiple comparisons. Active clusters are displayed on the MNI-2mm standard brain. BOTTOM – Bar graphs of the mean BOLD activation (y-axis) from significant clusters identified in the whole-brain interaction (see TOP). The graphs allow for qualitative visualization of the interaction effect.

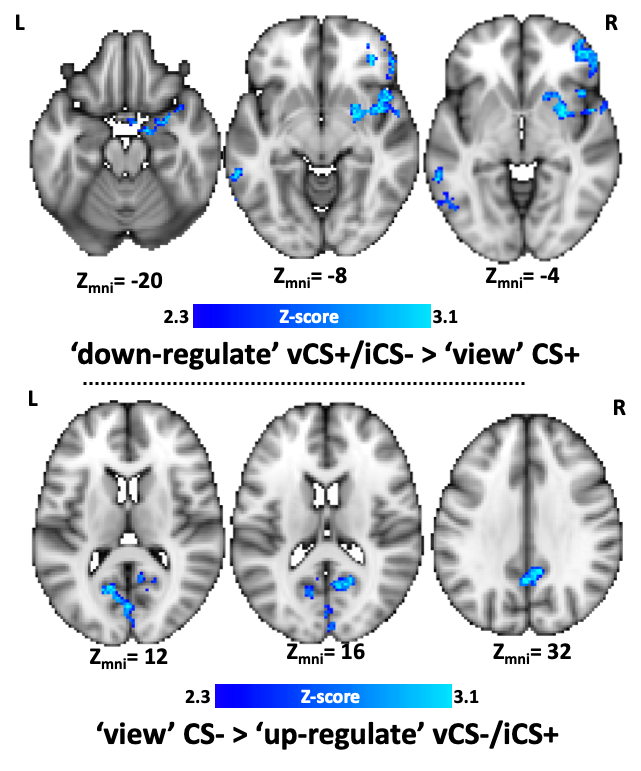

FIGURE S9. Experiment 2 Psycho-physiological interaction (PPI) whole-brain analysis. The seed for the PPI was created by taking the right aIn cluster produced by the spatial conjunction (see Figure S4) and excluding any voxels that were not in the insula (defined as the Harvard-Oxford right insula map). TOP – When down-regulating the CS+ (vCS+/iCS-) compared to simply viewing the CS+ there was greater functional connectivity between the right aIN and several regions including aspects right amygdala, right inferior frontal gyrus, and the striatum. BOTTOM – When simply viewing the CS- compared to up-regulating (vCS-/iCS+) there was greater functional connectivity between the right aIN and several regions including bilateral aspects of the visual cortex.
